# Distinct adaptation and epidemiological success of different genotypes within *Salmonella enterica* serovar Dublin

**DOI:** 10.1101/2024.07.30.605691

**Authors:** Cheryll M Sia, Rebecca L Ambrose, Mary Valcanis, Patiyan Andersson, Susan A Ballard, Benjamin P Howden, Deborah A Williamson, Jaclyn S Pearson, Danielle J Ingle

## Abstract

*Salmonella* Dublin is a host-adapted, invasive non-typhoidal *Salmonella* (iNTS) serovar that causes bloodstream infections in humans and demonstrates increasing prevalence of antimicrobial resistance (AMR). Using a global dataset of 1,303 genomes, coupled with *in vitro* assays, we examined the evolutionary, resistance, and virulence characteristics of *S*. Dublin. Our analysis revealed strong geographic associations between AMR profiles and plasmid types, with highly resistant isolates confined predominantly to North America, linked to IncC plasmids co-encoding AMR and heavy metal resistance. By contrast, Australian isolates were largely antimicrobial-susceptible, reflecting differing AMR pressures. We identified two phylogenetically distinct Australian lineages, ST10 and ST74, with a small number of ST10 isolates harbouring a novel hybrid plasmid encoding both AMR and mercuric resistance. Whereas the ST10 lineage remains globally dominant, the ST74 lineage was less prevalent. ST74 exhibited unique genomic features including a larger pan genome compared to ST10 and the absence of key virulence loci including SPI-19 which encodes a type VI secretion system (T6SS). Despite these genomic differences, the ST74 lineage displayed enhanced intracellular replication in human macrophages and induced less pro-inflammatory responses compared with ST10, suggesting alternative virulence strategies that may support systemic dissemination of ST74. The Vi antigen was absent in all ST10 and ST74 genomes, highlighting challenges for serotyping and vaccine development, and has implications for current diagnostic and control strategies for *S.* Dublin infections. Collectively, this study represents the most comprehensive investigation of *S*. Dublin to date and importantly, has revealed distinct adaptations of two genotypes within the same serovar, leading to different epidemiological success. The regional emergence and evolution of distinct *S.* Dublin lineages highlights the need to understand the divergence of intra-serovar virulence mechanisms which may impact the development of effective control measures against this important global pathogen.

## INTRODUCTION

Invasive nontyphoidal *Salmonella* (iNTS) infections are a major cause of global morbidity and mortality (1). Of the >2,500 NTS serovars, iNTS infections are associated with a relatively limited number of serovars. These include *Salmonella enterica* serovar Dublin (*S.* Dublin), *S.* Panama, *S.* Virchow, *S.* Choleraesuis and *S.* Typhimurium ST313 (2–4). *S.* Dublin, a bovine-adapted pathogen, is one of the leading iNTS serovars both in Australia and globally (5–7), and is associated with bacteraemia in a high proportion of infections (2,5–7). Symptomatic cattle may develop septicaemia and transplacental transmission (7,8), while asymptomatic infections in cattle enable persistence within the herd population, facilitating disease spread (7,9). *S*. Dublin causes zoonotic infections predominately through the consumption of contaminated dairy and meat produce (10–12), and as such is considered a ‘One Health’ pathogen. (10–12).

Antimicrobial resistance (AMR) in NTS has increased over the past several decades, leading to the reduction of therapeutic options (6,13). Although *S.* Dublin isolates are largely susceptible to therapeutics, AMR has become increasingly prevalent in *S.* Dublin (7,12,14–16). For example, a multidrug-resistant (MDR; defined as resistance to ≥ 3 antimicrobial classes (17)) lineage has recently been described in North America, characterised by varying profiles of AMR to ampicillin, streptomycin, chloramphenicol, sulphonamides, tetracyclines and third generation cephalosporins. These MDR *S.* Dublin isolates all type as sequence type 10 (ST10), and the AMR determinants have been demonstrated to be carried on an IncC plasmid that has recombined with a virulence plasmid encoding the *spvRABCD* operon (7,12,16,18). This has resulted in hybrid virulence and AMR plasmids circulating in North America including a 329kb megaplasmid with IncX1, IncFIA, IncFIB and IncFII replicons (isolate CVM22429, NCBI accession CP032397.1) (12,16) and a smaller hybrid plasmid 172,265 bases in size with an IncX1 replicon (isolate N13-01125, NCBI accession KX815983.1) (18).

*S.* Dublin is closely related to *S.* Enteritidis and *S.* Gallinarum (19), with differences in accessory genome content reflective of adaptation to different ecological niches (19). The majority of the STs reported for *S.* Dublin are ST10 or single-locus variants of this ST, although there is a second lineage, ST74, that is distinct from the main *S.* Dublin population (20). This uncommon *S.* Dublin lineage, ST74, may be variably serotyped as *S.* Dublin or *S.* Enteritidis dependent upon the strains used to generate typing sera in different laboratories (20), which may have also contributed to it not being identified. The antigenic formula of *S.* Dublin (1,9,12:g,p:-) is highly similar to *S.* Enteritidis (1,9,12:g,m:-), with three non-synonymous point mutations in *fliC* (Ala220Val, Thr315Ilc and Thr318Ala) differentiating the two serovars. Although the Vi capsular antigen (encoded by the *viaB* locus) is part of the antigenic definition for *S*. Dublin, it has been shown that the Vi antigen is absent in many *S.* Dublin isolates (20,21). Other putative virulence determinants in *S.* Dublin include *Salmonella* pathogenicity islands (SPIs) 6 and 19, which both encode type VI secretion system (T6SS) machineries (22,23). The presence of two T6SSs in *S*. Dublin contrasts with other NTS serovars, which encode a single T6SS (22,23). T6SSs are molecular nanomachines that deliver effector proteins into eukaryotic and/or prokaryotic cells to promote pathogen survival within multispecies and complex immune environments such as the gut. Recent studies have demonstrated a role for SPI-6 and SPI-19 in interbacterial competition by specific *S.* Dublin strains (24,25), however their effect on host immune processes remains largely unknown.

We previously undertook a genomic investigation of another emerging NTS lineage in Australia, *Salmonella enterica* serovar 4,[5],12:i:-, and identified several bacterial traits that may have contributed to the successful spread of this lineage (26–30). In particular, we hypothesised that flagellin deletion and MDR status may have contributed to host immune evasion and positive selection, respectively, in *Salmonella enterica* serovar 4,[5],12:i:- (31). Here, we sought to understand the contribution of virulence and resistance to the recent emergence of MDR *S*. Dublin. We interrogated a global dataset of 1,303 isolates collected from 13 countries on five continents and identified distinct geographical lineages associated with specific AMR and virulence profiles, including a lineage unique with increased capability for intracellular survival and immune evasion in human macrophages.

## RESULTS

### Distinct S. Dublin populations circulate globally

A total of 1,303 *S*. Dublin genomes were included in our analysis, spanning four decades (Figure 1A, Supplementary Tables 1-2). This dataset included 53 genomes from humans in Australia, and 1,250 genomes from Africa, Europe, North and South America collected from humans (*n*=462), animals (*n*=551), food (*n*=223) and other sources that were not stated (*n*=67) (Figure 1B, Supplementary Tables 1-2). In total, 1,235 genomes (99.4%) were ST10 and 58 were a single locus variant (SLV) of ST10, with two double locus variant genomes (e.g. ST4100) restricted to isolates from South America. The exceptions were ST74 and SLV ST1545 that were only detected in Australia (Figure 1C-E, Supplementary Table 3). All isolates, including the ST74 lineage, were serotyped *in silico* as *S.* Dublin. However, the ST74 lineage and a single ST10 isolate had codon profiles in *fliC* of Gly60, Leu138, Ala220, Thr315 and Ala318, in contrast to the remaining ST10 and SLV, which both had a profile of Gly60, Leu138, Val220, Ile315 and Ala318 (the previously described ‘Du2’ and ‘Du1’ electrophoretic profiles (21), respectively) (Supplementary Table 4).

**Figure 1.**
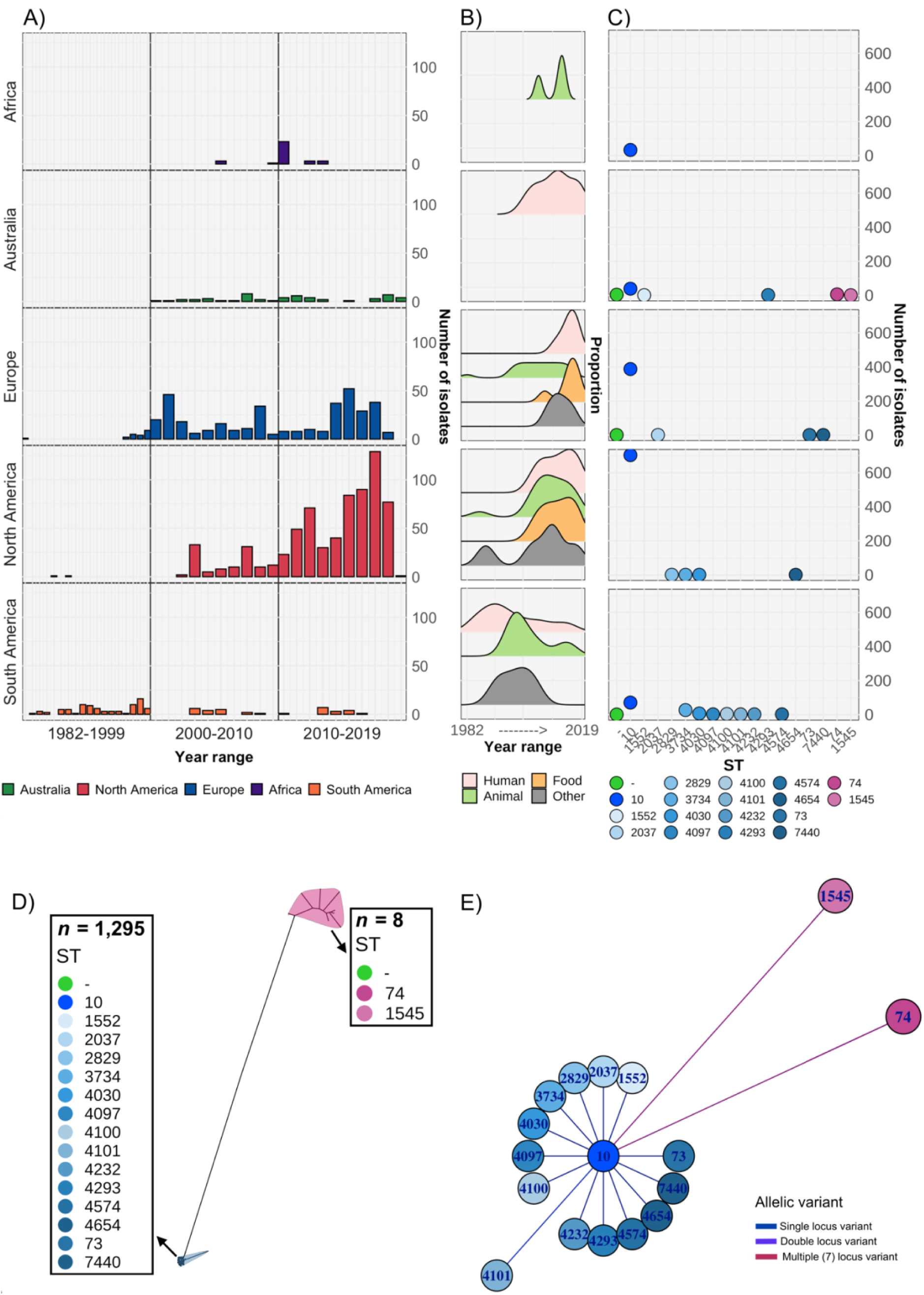
Overview of *S.* Dublin genomes in study. Geographical distribution and clonal relationship of 1,303 *S.* Dublin genomes in the study. A) Number of *S*. Dublin genomes according to geographical region and time interval of collection. B) Distribution of human, animal and food *S*. Dublin genomes collected over time. C) Distribution of *S*. Dublin sequence types (ST) from each region. Each coloured circle correlates to an ST. Circles in green indicate no ST was identified. D) Unrooted phylogenetic tree of 1,303 *S.* Dublin genomes. Two main lineages of ST74 (*n*=8) and ST10 (*n*=1,295) shown in blue and pink respectively. E) Relatedness of known sequence types (STs) to ST10. The nodes are labelled with the ST and the branch lengths are indicative of the number of dissimilar locus variants to ST10.

A maximum likelihood (ML) phylogeny was inferred from the 1,303 genomes, revealing two major lineages of *S.* Dublin, ST74 and ST10 (Figure 1D, Supplementary Figure 1). Within the ST10 lineage, five clades were identified using Bayesian Analysis of Population Structure (BAPS), inferred from the alignment of 2,957 core single nucleotide polymorphisms (SNPs). Four clades were broadly associated with geographical region (Supplementary Figure 1, Supplementary Table 1). The Australian ST10 isolates (*n* = 39) clustered in clade 1, while isolates from North America were associated with clade 5 (Supplementary Figure 1). The BAPS clade 2 were the isolates that didn’t have sufficient evidence to be split into further clades, hence are spread across the tree spanning a range of geographical regions.

### Emergence of S. Dublin lineages in the 20^th^ century

To investigate the emergence of *S*. Dublin clades, Bayesian phylodynamic analysis was undertaken on a representative subset of 660 genomes, selected for phylogenetic, geographical and AMR diversity (see methods). We inferred a maximum clade credibility (MCC) phylogeny from an alignment of 3,971 SNPs using the highest supported model of a relaxed lognormal molecular clock and coalescent exponential tree prior. The estimated substitution rate was 1.05 x 10^−7^ (95% highest posterior density (HPD): 1.16 x 10^−7^– 9.41 x 10^−8^), slightly higher than previous estimated mean substitution rates for host-restricted *Salmonella* serovars (*S*. Typhi and *S*. Paratyphi A) but lower than rates reported for host-generalist NTS serovars *S*. Kentucky and *S*. Agona (32).

The topology of the MCC tree was consistent with the ML tree, with geographically associated sub-clades were observed in the MCC tree (Figure 2, interactive tree is available in Microreact under https://microreact.org/project/sdublinpaper). The most recent common ancestor (MRCA) for the *S.* Dublin global population was inferred to be 1901 (HPD 1878-1924), with the MRCA of the different clades and sub-clades largely falling between 1950 and 2000. For example, the most recent common ancestor (MRCA) of clade 5 MDR isolates from North America was estimated to be ∼1964 (HPD 1952-1974) (Figure 2). The MRCA of the Australian sub-clade was estimated to be ∼1967 (HPD 1953-1980); with the Australian and North American isolates inferred to have a MRCA ∼1931 (HPD 1914-1947). Moreover, three clades (1, 3, and 4) had European sub-clades associated with Sweden, the UK, and Denmark, respectively that were all largely susceptible to antimicrobials. Some regions, such as Europe and South America, were associated with several sub-clades of *S.* Dublin, potentially suggesting multiple importations and expansions, while other regions, such as Australia were comprised of a single sub-clade within clade 1.

**Figure 2.**
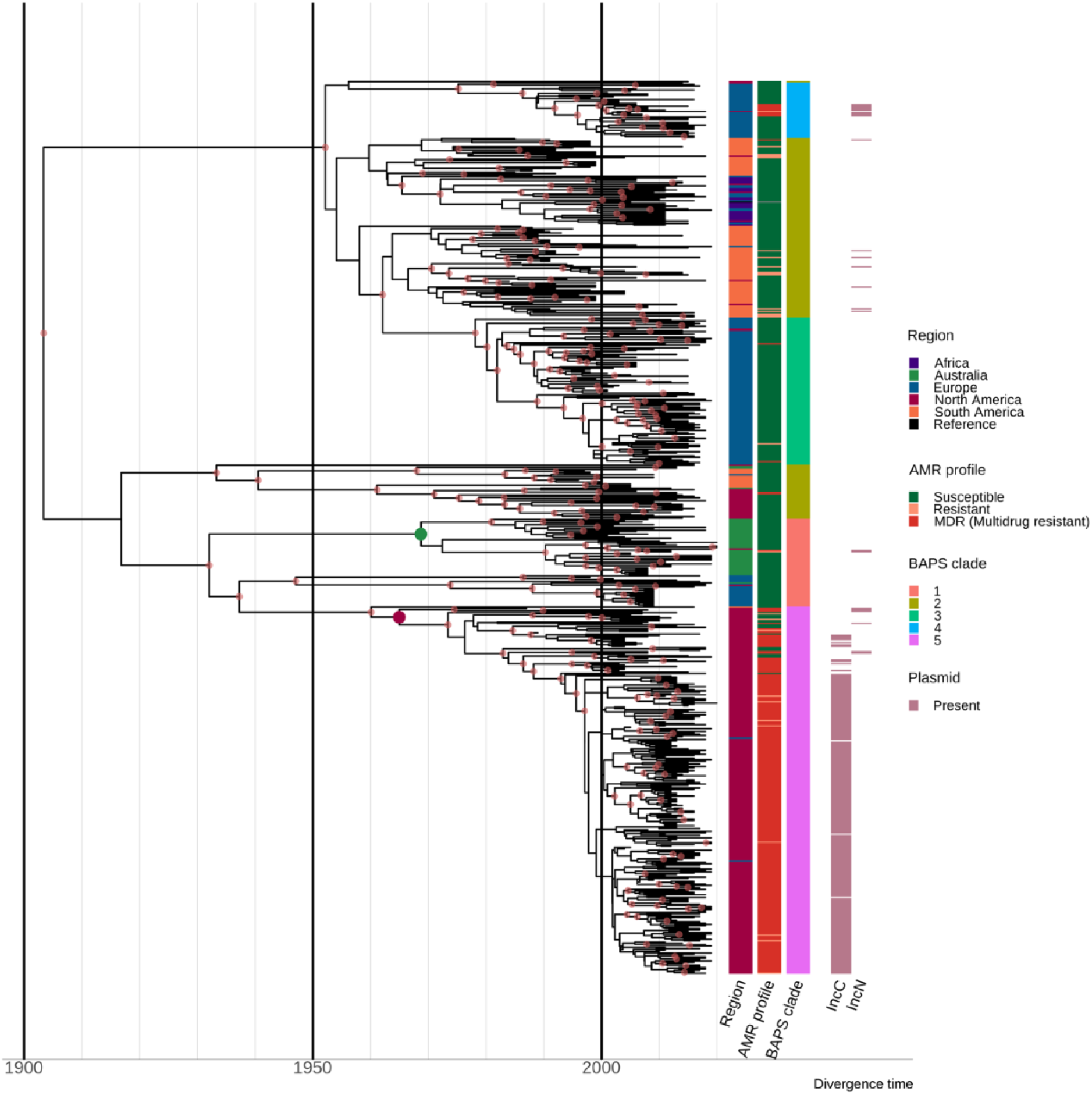
Estimation of divergence time. Maximum clade credibility (MCC) tree of the 667 *S.* Dublin ST10 genomes. The time in years is shown on the x-axis. From left to right, the coloured heatmaps refer to the region of origin, the susceptibility profile, the BAPs clade, and the presence of IncC and IncN plasmid types. The green dot denotes the most recent common ancestor (MRCA) of the Australian subclade, while the maroon dot represents the MRCA of clade 5 MDR isolates from North America. Internal nodes coloured chestnut red have a posterior probability ≥ 0.95.

### Resistance determinants are associated with geographical regions

Different plasmid types co-occurred with different AMR and HMR determinants in the *S.* Dublin population, and these were also associated with differential geographical distribution (Figure 2, Supplementary Figure 3-4, Supplementary Table 1). Overall, 51.5% (667/1,295) of *S.* Dublin genomes had at least one AMR determinant, with 632/667 (94.8%) isolates classified as MDR. MDR *S.* Dublin was predominantly identified from North American isolates, although some AMR determinants were detected in *S.* Dublin from other geographic regions (Figure 3, Supplementary Figures 2-3). AMR *S.* Dublin collected from animals was common (*n*=305/667; 45.7%). AMR in *S.* Dublin from food sources and humans was similar in proportion (*n*=174/667; 26.1% and *n*=173/667; 25.9% respectively). Although these profiles are likely impacted by the sampling frame of different studies, the source of collection was not considered as a selection criterion for this study, and as such, may broadly represent the AMR profiles of *S.* Dublin from different sources (Figure 3A-B, Supplementary Figure 1).

**Figure 3.**
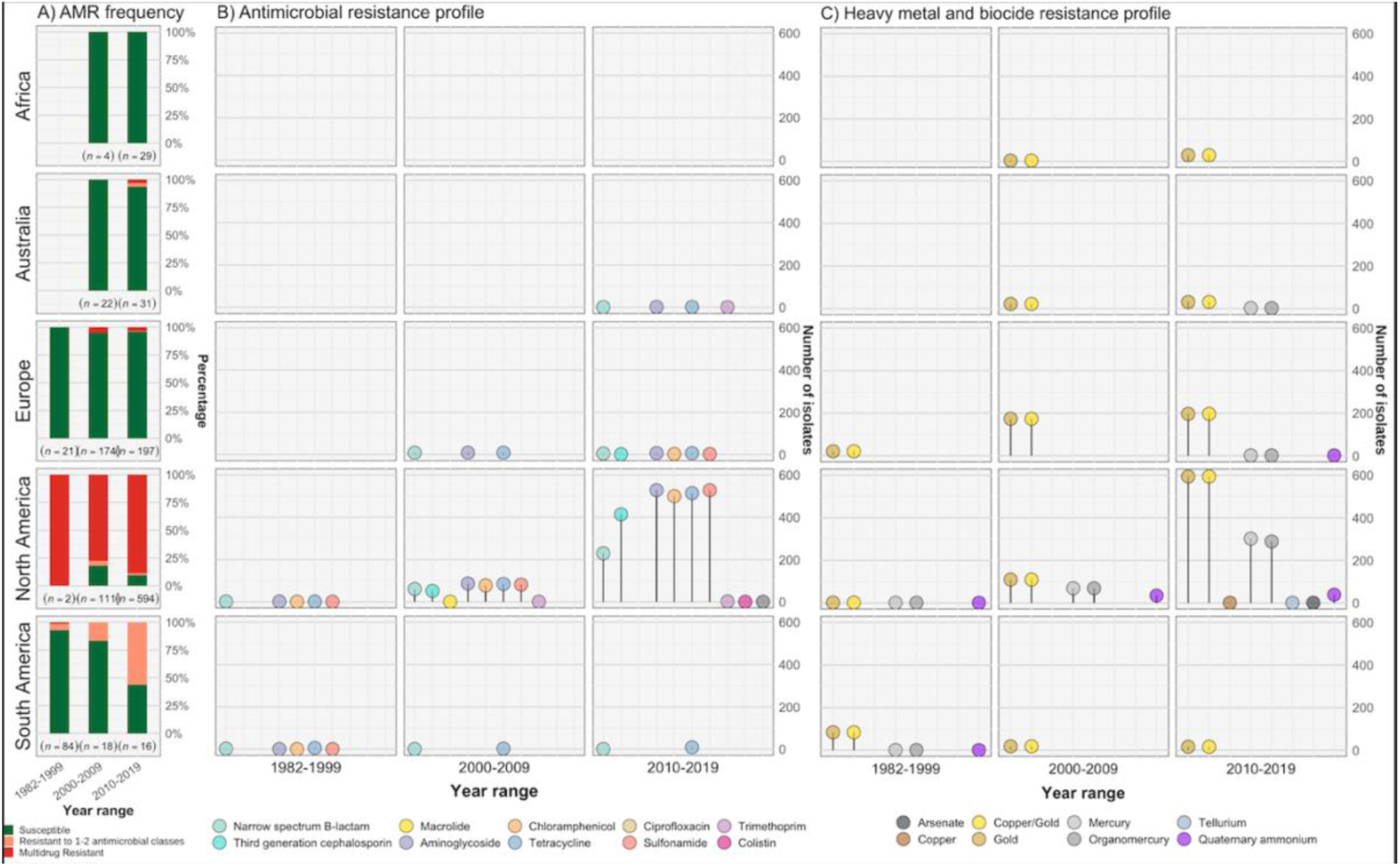
Resistance profiles of *S.* Dublin genomes. Observed resistance profiles of *S*. Dublin genomes spanning three-time intervals for each region. A) Frequency of susceptible and resistant *S*. Dublin genomes from each time period B) Antimicrobial resistance profiles observed spanning three time periods. The colour of the circle corresponds to an antimicrobial class, while the height is relative to the number of genomes. C) Heavy metal and biocide resistance profiles observed. The colour of the circle corresponds to its respective heavy metal/biocide class, while the height is relative to the number of genomes.

The most prevalent AMR determinants were *strA*-*B* (*n*=620), *floR* (*n*=556), *tet*(A) (*n*=608) and *sul2* (*n*=596) (Supplementary Figure 2). AMR mechanisms to ciprofloxacin (triple point mutations in quinolone resistance determining regions *gyrA*[83:S-F]*, gyrA*[87:D-N]*, parC*[80:S-I]), macrolides (*mph*(A)), and colistin (*mcr-9.1*) were each detected in single genomes from *S.* Dublin clade 5 at a low frequency of 0.08% (*n*=1/1,303) in the dataset. The only third generation cephalosporin (3GC) resistance mechanism detected was *bla*_CMY-2_, present in 467/667 (70%) of AMR *S.* Dublin genomes. Although *strA*-*B*, *tet*(A) and *bla*_TEM-1_ were observed in multiple clades, *floR*, *sul2* and *bla*_CMY-2_ were restricted to North American isolates that fell in clade 5 (Supplementary Figure 2-3). This MDR profile, in addition to *strA-strB* and *tet*(A), was mediated by IncC plasmids in 593 genomes (*n*=509 from the United States of America [USA], *n*=82 from Canada and *n*=2 from the UK) (Supplementary Figure 3, 4C). Nearly all clade 3 isolates linked to the UK had no AMR determinants, except for four isolates collected between 2010-2015 (Supplementary Figure 1-3, Figure 3, Supplementary Table 1).

In addition to AMR determinants, we characterised biocide and heavy metal resistance (HMR) profiles of the *S.* Dublin isolates. Gold resistance determinants *golS* and *golT* were ubiquitous in *S.* Dublin (*n*=1,302/1,303; 99.9%) (Figure 3C, Supplementary Figures 2-3). Consistent with previous findings biocide and HMR determinants, including *qacEdelta1* and mercury resistance genes (*merABDEPRT*), commonly co-occurred with AMR determinants. Notably, the *mer* operon was frequently associated with the IncC MDR plasmid (90.1%; *n*=534/593) (Supplementary Figure 3, Supplementary Table 1).

### Characterisation of S. Dublin plasmids and identification of a novel hybrid plasmid

Both IncX1 (*n*=1269/1303; 97.4%) and IncFII(S) (*n*=1213/1303; 93.1%) plasmid replicons were found at high frequency within the dataset, in line with previous findings (12). Three plasmid replicons, namely IncN, IncX1 and IncFII(S), and a MOBF/MOBP relaxase, were detected in one of the Australian *S*. Dublin ST10 genomes from clade 1 (AUSMDU00035676). Comparative analysis revealed the AUSMDU00035676 plasmid lacked the *spv* operon and had low homology to reference IncN and IncX1/IncFII(S) plasmids previously associated with *Salmonella* (Supplementary Figure 5, Supplementary Table 5). The loss of the *spv* operon may be associated with a less invasive phenotype (33). This finding suggests the detection of a potentially novel hybrid plasmid with IncN/IncX1/IncFII(S) markers that encodes AMR and HMR determinants flanked by IS26 elements, which may aid gene mobilisation (34). In contrast, the AUSMDU00056868 plasmid had IncX1/IncFII(S) plasmid replicon markers and was found to not encode any mechanisms for resistance. This plasmid was >99% homologous to known virulence reference *S.* Dublin plasmids pOU1115 and pSDU2_USMARC_69807 and encoded the *spv* operon (Supplementary Table 5, Supplementary Figure 6). This suggests that the *S.* Dublin virulence plasmid is conserved, while the *spv* operon that has been shown to enhance disease severity (14,16,35–37) is maintained on the plasmid. Complete annotations of both AUSMDU00035676 and AUSMDU00056868 can be found in Supplementary Tables 6 and 7, respectively.

### Diversity in virulence determinants across S. Dublin lineages

To investigate the potential for invasiveness within *S.* Dublin, we applied a previously described bioinformatic tool to our dataset of 1,303 *S.* Dublin genomes (38). In brief, this tool infers the likelihood of invasiveness by assigning an index score between 0 to 1 that reflects the level of genome degradation, whereby a higher score suggests genomic features that may be more associated with a narrower host range and great likelihood of invasiveness (38). We performed initial validation work using 36 complete publicly available reference genomes, covering known invasive and non-invasive phenotypes (Supplementary Table 8). Using this validation set, invasive and largely host-restricted serovars, including *S*. Choleraesuis, *S*. Paratyphi and *S*. Typhi, scored >0.5, suggestive of an invasive genomic profile (Figure 4A, Supplementary Table 8). Conversely, non-invasive NTS serovars with broader ecological niches, including *S*. Anatum, ST19 *S*. Typhimurium and *S.* Derby, scored <0.5 (Figure 4A, Supplementary Table 8). When applied to our *S.* Dublin dataset, only eleven genomes (*n*=11/1,303; 0.84%) had an index score < 0.5. These included three ST10 genomes and all eight ST74 genomes (Figure 4B, Supplementary Table 9). Based on descriptive analysis, no apparent relationship was observed between index scores and region or source attribution (Figure 4B).

**Figure 4.**
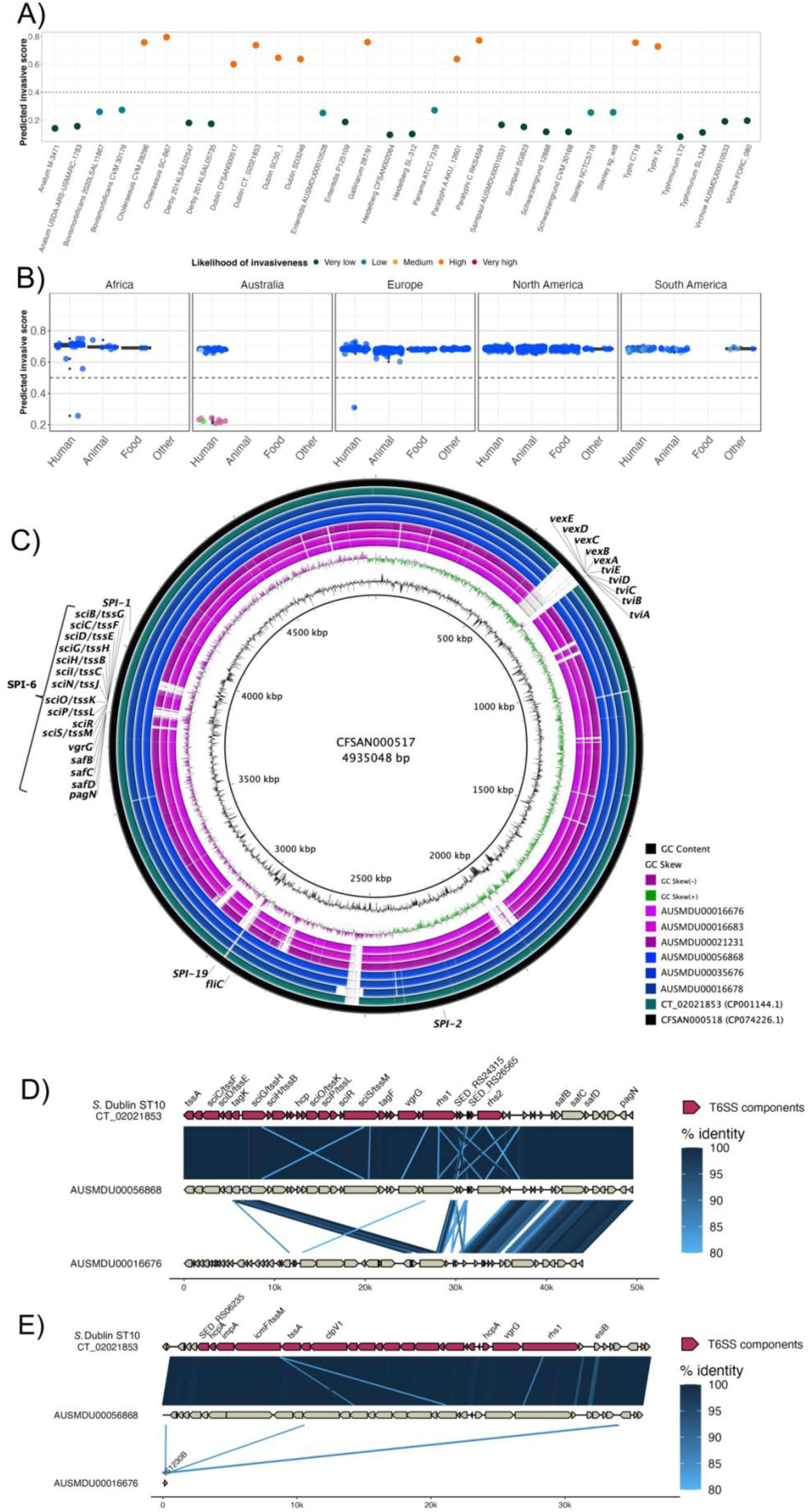
Invasive factors associated with *S.* Dublin genomes. Correlation of *Salmonella* pathogenicity islands (SPIs) to *S*. Dublin’s invasiveness. A) Validation of the likelihood of invasiveness prediction tool using a variety of publicly available genomes consisting of non-invasive and invasive *Salmonella* serovars (Supplementary Table 8). B) The correlation of index scores to the source and its sequence type based on the region. * indicates a statistically significant p-value between the index scores of two sources. C) Chromosomal assemblies of ST10 (blue) and ST74 (pink) genomes aligned to *S*. Dublin chromosomal reference CT_02021853 (CP001144.1) with a lower limit of 80% identity D) Aligned sequence of SPI-6 region of an ST10, AUSMDU00056868, and an ST74, AUSMDU00016676, in comparison to SPI-6 region of reference CT_02021853. E) Aligned sequence of SPI-19 region of an ST10, AUSMDU00056868, and an ST74, AUSMDU00016676 in comparison to SPI-19 region of reference CT_02021853. SPI, *Salmonella* pathogenicity island.

Putative virulence determinants identified in *S*. Dublin genomes included SPIs, the *spv* operon (16), prophages and other individual genes (Supplementary Table 9). Some putative virulence factors, such as *pagN* [cell adhesion and invasion (39,40)] and *pgtE,* [virulence (27,41)], were found in 1,303 (100%, *n*=1,303/1,303) and 1,302 (99.9%, *n*=1,302/1,303) genomes respectively, and no SNPs in the promoter region of *pgtE* were detected, a mutation previously reported in invasive *S*. Typhimurium ST313 (27). In contrast, ST313-gi of unknown function (27,41), the Gifsy-2 like prophage [unknown function (27,41)] and *ggt* gene thought to be involved in gut colonisation (35), were detected in the major ST10 *S.* Dublin lineages, and absent in ST74 *S.* Dublin. The *viaB* operon encoding the Vi capsular antigen was only detected in SLV ST73, a historical isolate from France that displayed the Du3 electrophoretic type that was consistent with a previous study (21). This suggests that the Vi antigen is not conserved in the *S.* Dublin population or associated with virulence.

The presence of SPI-6 and SPI-19 differed across *S.* Dublin isolates. Both SPIs encode T6SSs that provide a means for injecting effector proteins into adjacent competing bacteria or host cells for immune evasion (23,42). SPI-19 was absent in the ST74 *S.* Dublin lineage but present in all ST10 *S.* Dublin (Supplementary Figure 7). The genetic differences between ST74 and ST10 were reflected in the disparity of their index scores of <0.5 and >0.5, respectively. The SPI-6 region was detected in all isolates, however <90% sequence coverage was detected in 422/1,303 genomes (32.4%). Long read sequencing on three genomes of the ST74 lineage and three genomes from the ST10 clade 1 lineage corroborated these findings (Figure 4C-E), demonstrating truncation of SPI-6 with the T6SS machinery (Figure 4D). Long-read sequencing also revealed an absence of SPI-19 in the three ST74 isolates and presence of SPI-19 in the three ST10 isolates (Figure 4C, E). The absence of SPI-19 in ST74 suggests that this deletion event occurred during its divergence from the primary *S*. Dublin lineage, ST10.

### Infection of human macrophages reveals distinct intracellular replication patterns between ST10 and ST74 lineages

Given the distinct genomic profiles of ST10 and ST74 lineages, we compared the phenotypic profile of a selection of Australian isolates from each lineage *in vitro*. The immortalised human macrophage cell line, THP-1, was used to assess invasion, intracellular replication and host cytotoxicity induced by a subset of clinical isolates from the two lineages. Specifically, we assayed an ST10 population comprising eleven isolates, including an isolate from the highly related ST4293 lineage, and an ST74 population comprising seven isolates including an isolate from the highly related ST1545 lineage (Supplementary Table 10). Although predicted invasive index scores were higher for the ST10 population (Figure 4B), each isolate failed to replicate more than 2-fold over 24 hours in THP-1 cells (Figure 5A), whereas isolates within the ST74 population displayed a sustained increase in fold change of intracellular replication, up to 8-fold, over 24 hours (Figure 5B). Comparing bacterial growth (CFU/ml) across both sequence types, both lineages were comparable for initial bacterial uptake (Figure 5C) however, the ST74 population replicated to significantly higher levels in THP-1 macrophages at 9- and 24-hours post-infection (hpi) compared to the ST10 population (Figure 5D-F). Given that NTS also replicate in epithelial cells, we utilised an immortalised human intestinal epithelial cell line, HT-29, to assess *S.* Dublin replication. In contrast to the results obtained in macrophages, we found that 6 out of 11 isolates within the ST10 population replicated more efficiently (up to 10-fold) in HT-29 cells, whereas all seven isolates within the ST74 population showed minimal replication (up to 2-fold) over 24 hours (Supplementary Figure 8A-C).

**Figure 5.**
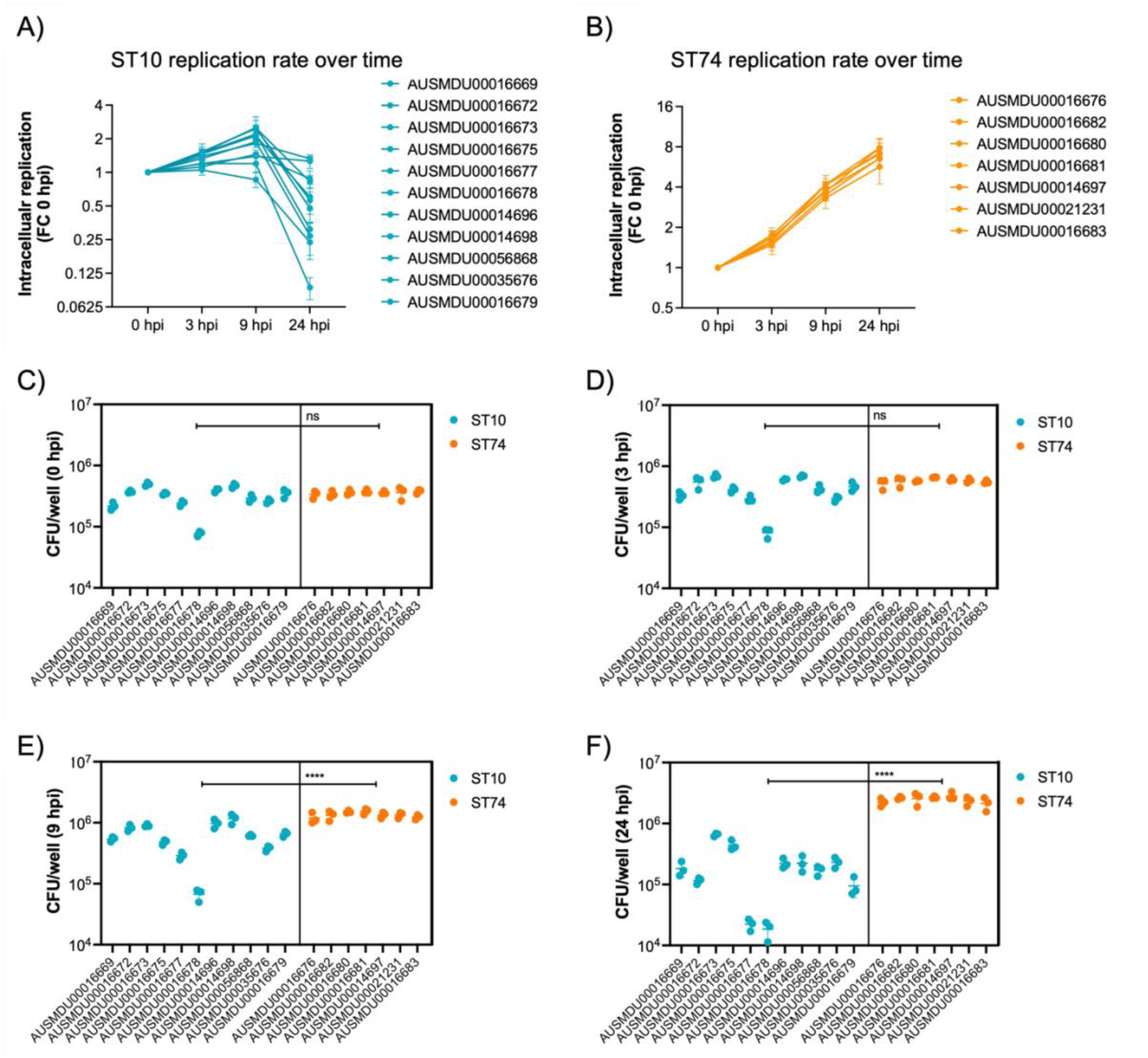
*S.* Dublin ST74 isolates demonstrate increased intracellular replication in human macrophages over ST10 isolates. Differentiated THP-1 (human monocyte) cells were infected at an MOI:10 with selected ST10 and ST74 *S.* Dublin isolates. **A, B)** THP-1 cells were lysed, and intracellular bacteria enumerated as fold change in replication and **C-F)** CFU/well at time 0, 3, 9 and 24 hpi. For **A)**, each measurement at times 3, 9 and 24 hpi represents the fold change in CFU/well compared to 0 hpi, the start of infection, recorded as a biological replicate (performed in technical triplicate), with error bars indicating ± 1 standard deviation of *n* = 3 biological replicates. For **C-D**), each dot represents CFU/well of a biological replicate (performed in technical triplicate), with error bars indicating ± 1 standard deviation of *n* = 3 biological replicates. Statistical significance was determined by nested student’s t-test. MOI, multiplicity of infection, hpi, hours post-infection, CFU, colony-forming units, FC, fold-change.

### Increased replication of ST74 isolates in human macrophages does not correlate with increased host cell death

It is well documented that protype *S*. Typhimurium (ST19) potently induce macrophage cytotoxicity both *in vitro* and *in vivo* (43). Here, we utilised a cellular lactate dehydrogenase (LDH) release assay to assess THP-1 cytotoxicity in response to infection with *S.* Dublin ST74 and ST10 populations. Surprisingly, the substantial intracellular replication of the ST74 population in THP-1 cells induced significantly less host cell cytotoxicity at 3 and 9 hpi compared with the ST10 population suggesting potential immune evasion by ST74 (Figure 6A, B). By 24 hpi however, cytotoxicity induced by both ST74 and ST10 populations was relatively high, with no significant difference observed across the sequence types (Figure 6C). In contrast, there was minimal cytotoxicity observed in infected HT-29 cells across the board (Supplementary Figure 8D), suggesting a less potent activation of host cell death in infected epithelial cells compared with macrophages.

**Figure 6.**
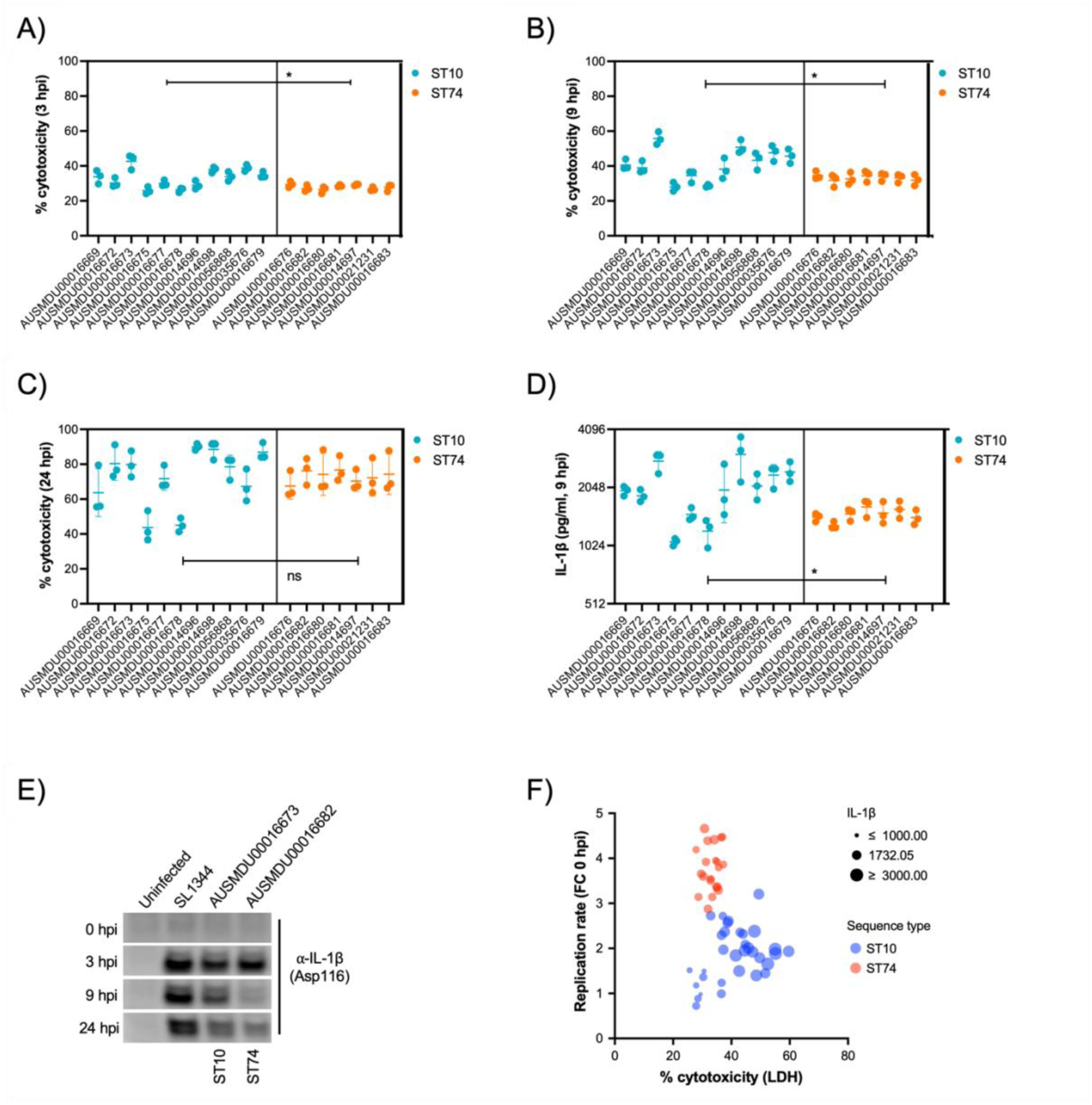
*S.* Dublin ST74 isolates induce comparable cytotoxicity to ST10 isolates in human macrophages, despite increased intracellular growth. **A-C)** Cell supernatants were assayed for LDH as a measure of cytopathic effects of infection, and % cytotoxicity was calculated by comparison with 100% lysed uninfected control cells. Samples were collected from THP-1 cells at 3, 9 and 24 hpi. Each point represents the % cytotoxicity of a biological replicate (performed in technical triplicate), with error bars indicating ± 1 standard deviation of *n* = 3 biological replicates. Statistical significance was determined by nested student’s t-test. **D)** IL-1ß secretion from THP-1 cells infected for 9 hours with *S.* Dublin isolates. Error bars indicate ± 1 standard deviation of *n* = 3 biological replicates and statistical significance determined by student’s t-test **E)** Immunoblot of cleaved IL-1ß (Asp166) in THP-1 cell lysates infected with *S.* Typhimurium reference strain SL1344, *S.* Dublin ST10 (AUSMDU00016673), or *S.* Dublin ST74 (AUSMDU00016682) over 0, 3, 8 and 24 hours. **F)** Scatter plot comparing replication rate, % cytotoxicity and IL-1β secretion from THP-1 cells infected with *S.* Dublin isolates at 9 hpi. Each dot represents each biological replicate of three (averaged from three technical replicates). MOI, multiplicity of infection, hpi, hours post-infection, CFU, colony-forming units, LDH, lactate dehydrogenase.

To understand how the ST74 population may be supressing macrophage death at earlier stages of infection, we measured protein secretion of IL-1ß via cytokine release and immunoblot assays at 9 hpi, respectively. IL-1ß is a potent pro-inflammatory cytokine that is released upon activation of the NLRC4 inflammasome by components of the *Salmonella* type III secretion apparatus (44), resulting in a lytic cell death process known as pyroptosis (45). Here, we observed significantly less IL-1ß release by THP-1 cells infected with isolates from the ST74 population at 9 hpi compared with those from the ST10 population (Figure 6D), directly reflecting the LDH cytotoxicity data represented in Figure 6B. To further support our observations, we selected single representative isolates of ST74 and ST10 to perform immunoblot analyses of cleaved IL-1ß (activated form) within THP-1 cells. Here, ST74 induced less IL-1ß cleavage at 9 hpi compared to the ST10 (Figure 6E). We included the ubiquitous laboratory NTS reference strain, *S.* Typhimurium SL1344 in the immunoblot analysis as this is known to potently induce pyroptosis in human macrophages (46). As expected, SL1344 induced significant IL-1ß cleavage during infection of THP-1 macrophages (Figure 6E). To assess the correlation of cellular cytotoxicity as measured by LDH, IL-1ß release and intracellular replication, we generated a scatter plot of all isolates within the ST74 and ST10 populations. Here we can clearly demonstrate that isolates from the ST74 population have increased replicative ability, induce less cytotoxicity, and lower levels of IL-1ß release (Figure 6F) compared with those within the ST10 population. There were however, some ST10 isolates that induced similarly low levels of host cell death, presumably due to lower levels of intracellular replication (Figure 6F and Supplementary Figure 9A, B). These isolates generally clustered away from the ST10 isolates that induced more cell death (Supplementary Figure 9C).

A pangenome analysis of the isolates from both ST10 and ST74 lineages revealed that out of a total of 5,792 genes, there were 473 and 1,143 genes specific to the ST10 and ST74 lineage, respectively (Supplementary Figure 10, Supplementary Table 11). Further, out of 1,143 genes, there were 362 core genes present in all isolates from the ST74 lineage (Supplementary Table 11). To highlight these differences, an SNP distance matrix was generated, revealing a minimum of 2,402 SNPs between the ST10 and ST74 lineages (Supplementary Table 12). While these findings require further investigation, an increased gene content and genome size were consistently observed in all isolates of the ST74 lineage (Supplementary Tables 11 and 13). These results indicate a potential link between the unique genome content of the ST74 lineage and alterations in host responses and replicative ability.

## DISCUSSION

Our data provide comprehensive insights into the emergence and spread of *S*. Dublin, with distinct geographic distribution related to AMR and virulence. Analysing the largest global dataset of *S*. Dublin genomes to date, we show that highly resistant *S.* Dublin is almost entirely confined to North America, with emergence in the late 20^th^ century and ongoing local microevolution. Consistent with previous studies, the North American *S.* Dublin isolates in clade 5 were characterised by the presence of an IncC plasmid, often with co-location of AMR and HMR determinants (16,18), while the clades circulating in Europe were associated with IncN plasmids (47). Previous studies have explored these IncC plasmids in detail. The use of biocides and heavy metals in the agri-food industry may facilitate co-selection, contributing to the persistence of AMR (48). This highlights the potential for selection pressure in one ecological niche (e.g. the use of biocides in farming or heavy metals in animal feeds) leading to AMR in another (e.g. AMR in humans).

The phylogeographic separation, AMR and plasmid profiles of *S.* Dublin lineages is consistent with other studies (7,12,49–53), and is likely to be influenced by several factors, including local AMR use and patterns of livestock movement. Given the livestock reservoir of *S.* Dublin is mostly host-restricted to cattle, it is conceivable that both historic and contemporary global cattle movement may have contributed to region-specific differences in *S.* Dublin. For example, Australian *S.* Dublin genomes in clade 1 shared a common ancestor with North American isolates in clade 5. During the early 1930’s, Brahman cattle were imported from the U.S into Australia (54), supporting our phylodynamic finding that the common ancestral link between *S*. Dublin isolates from North America and Australia emerged in the early-mid 20^th^ century. Following region-specific introductions, subsequent local antimicrobial and husbandry practices may have contributed to the local evolution of *S.* Dublin (50,55).

Of note, among *S.* Dublin isolates from Australia, genomic analysis (cf. serotyping) revealed a smaller population of ST74 isolates. *S.* Dublin ST74 was previously identified as *S*. Enteritidis ST74, and sits as an intermediate between *S*. Dublin and *S*. Enteritidis (20), and has been understudied to date in studies of *S.* Dublin. The difference between the antigenic formula of *S*. Dublin and *S*. Enteritidis is in the H1 flagella region, which is typed as “*g,p*” for *S.* Dublin and “*g,m*” for *S.* Enteritidis. ST74 *S.* Dublin genomes were characterised by: (i) a lack of any AMR or HMR determinants; (ii) lack of complete SPI-6 regions, specifically the T6SS, and (iii) a lack of putative virulence determinants including ST313-gi, Gifsy-2 like prophage and SPI-19, which encodes a T6SS mechanism for intestinal colonisation and survival that interacts with the host by mediating macrophage cytotoxicity (55).

The lack of the Vi antigen in 1,302/1,303 (99.9%) *S*. Dublin genomes has several implications. First, the antigenic formula of *S*. Dublin, 1,9,12[Vi] may need to be reviewed if most *S.* Dublin genomes lack the machinery to express the Vi capsule. These limitations with serotyping have been highlighted in a previous study, which suggested a transition to a novel genotyping scheme that would prevent these phenotypic discrepancies (56). Second, the Vi antigen is a vaccine target for *S.* Typhi (57), and the Vi antigen has been explored as a target for *S.* Dublin (58). While previous studies have noted that some *S.* Dublin genomes may be capable of producing the Vi antigen (58), the genomic data from this study shows that these genomes are in the minority of the *S.* Dublin population and may not be stable targets for *S.* Dublin specific vaccines.

Our phenotypic data demonstrated a striking difference in replication dynamics between ST10 and ST74 populations in human macrophages. ST74 isolates replicated significantly over 24 hours, whereas ST10 isolates were rapidly cleared after 9 hours of infection. ST74 induced significantly less host cell death during the early-mid stage of macrophage infection, supported by limited processing and release of IL-1ß at 9 hpi. While NTS are generally potent inflammasome activators (60), most data that support this observation have been generated using laboratory-adapted *S*. Typhimurium strains, often ST19. Our findings suggest that ST74 isolates may employ immune evasion mechanisms to avoid host recognition and activation of cell death signaling in the early stages of infection. Similar trends have been observed with other invasive *S. enterica* serovars, including *S*. Typhimurium ST313, which induces less inflammasome activation than ST19 during murine macrophage infection (61). This suppression of inflammatory cell death may facilitate increased replication and dissemination of ST74 isolates at later stages of infection. Consistent with this, we observed comparable cytotoxicity between ST10 and ST74 populations at 24 hpi, suggesting ST74 induces cell death via alternative mechanisms once intracellular bacterial numbers are unsustainable. Further research is needed to identify genomic factors underpinning these observations.

In this study, ST74 isolates associated with human disease (n=7) were identified from faecal samples (cases of gastroenteritis), whereas 18/40 ST10 isolated were from invasive cases of human disease (from blood or abscess). In line with this, our machine learning approach predicted lower invasiveness for ST74 compared to ST10. However, this does not align with what we observed *in vitro,* i.e., higher replication of ST74 in macrophages compared with ST10. We speculate that the increased genomic complexity of ST74 may support higher replication in macrophages and that increased intracellular replication may enhance systemic dissemination, though this would require robust *in vivo* validation. Invasiveness of *S*. *enterica* is often linked to genome degradation (4,62–64). However, this is mostly based on studies of human-adapted iNTS (ST313) and *S*. Typhi, leaving open the possibility that the additional genomic richness of ST74 supports survival in diverse host species. An uncharacterised virulence factor or single nucleotide polymorphism within virulence genes of ST74 may also explain this replication advantage. Interestingly, the absence of SPI-19 in ST74, which encodes a T6SS, may reflect adaptation to enhanced replication in macrophages. SPI-19 has been linked to intestinal colonisation in poultry (23,56) and mucosal virulence in mice (56). It is possible that the efficient replication of ST74 in macrophages may compensate for the absence of SPI-19, relying instead on phagocyte uptake via M cells or dendritic cells. Collectively, these findings highlight phenotypic differences between *S*. Dublin populations ST10 and ST74, whereby enhanced intra-macrophage survival of ST74 could promote invasive disease, whereas the prevalence of ST10 may relate to better intestinal adaptation and enhanced faecal shedding. These findings highlight important knowledge gaps in zoonotic NTS host-pathogen interactions and drivers of emerging invasive NTS lineages with broad host ranges.

This study has limitations, including a focus on ST10 isolates from clade 1, which do not represent global phylogenetic diversity. Nonetheless, our pangenome analysis identified >900 uncharacterised genes unique to ST74, offering potential targets for future research. Another limitation is the geographic bias in available genomes, with underrepresentation from Asia and South America. This reflects broader disparities in genomic epidemiological studies but may improve as public health genomics capacity expands globally.

Overall, this study represents the most comprehensive genomic study of *S.* Dublin to date, systematically characterising antimicrobial, heavy metal and biocide resistance determinants, in addition to exploring the virulome in the global population of *S.* Dublin. Importantly, our phenotypic data reveals the caution required when making generalised statements about infection outcomes and virulence mechanisms of *Salmonella* serovars. Our data clearly shows that within the Dublin serovar, there are distinct phenotypic differences observed across sequence types and host cell types, all of which could impact significantly on future development of effective therapeutics or vaccine strategies for NTS infections. While the data did not indicate an increasing trend of iNTS associated with *S*. Dublin, the potential public health risk of this invasive pathogen suggests it may still warrant considering it a notifiable disease, similar to typhoid and paratyphoid fever.

## METHODS

### Setting and dataset

For Australian isolates, whole genome sequencing (WGS) was performed on all viable *S*. Dublin isolates from human clinical samples received at the Microbiological Diagnostic Unit Public Health Laboratory (MDU PHL), the primary bacterial reference laboratory for the State of Victoria in Australia. In total, 53 *S*. Dublin genomes between 2000-2019 were included. For publicly available data, we included *S*. Dublin sequences that had associated metadata (geographical origin and date of sample collection), providing a total of 1,250 publicly available sequences. Data were collected in accordance with the Victorian Public Health and Wellbeing Act 2008. Ethical approval was received from the University of Melbourne Human Research Ethics Committee (study number 1954615.3).

### Genome quality control, assembly and serovar prediction

Genomes were assembled and quality controlled using the Nullarbor pipeline v2.0.20191013 (https://github.com/tseemann/nullarbor). Briefly, Skesa v2.4.0 (59) was used to assemble the genomes before determining the sequence type (ST) using the multilocus sequence typing (MLST) scheme ‘*senterica’* within mlst v2.19.0 (https://github.com/tseemann/mlst). Kraken v1.1.1 (60) was then used to confirm the taxonomic classification. Snippy v4.6.0 (https://github.com/tseemann/snippy) with parameters set to a 0.90 variant rate proportion against a minimum coverage of 10X mapped the reads against a well-studied reference genome, *S*. Dublin CT_02021853 (NC_011205.1). To be included in subsequent analysis, genomes had to pass the following quality control parameters: mapped to ≥ 80% of the reference genome; had a ≥ 80% taxonomic sequence similarity to *S. enterica*; N50 >25000, and <300 contigs. To verify that each genome was *S*. Dublin, all sequences were typed using *Salmonella* in silico Typing Resource (SISTR) v1.1.1 (61). The electrophoretic type of each isolate was determined using Antimicrobial Resistance Identification by Assembly (ARIBA) v2.14.5 (62) against the reference Du 2 (Genbank accession: M84972.1). The draft genome assemblies were annotated using Prokka v1.14.6 (63).

### Core genome phylogenomic analysis

Variant calls from snippy were used to construct a core genome single nucleotide polymorphism (SNP) alignment using snippy-core v4.6.0 (https://github.com/tseemann/snippy). Phaster v2020-12-22 (64) was used to create a BED file containing prophage regions. The option ‘—mask’ in snippy-core was used to exclude the identified prophage regions from the BED file. Recombination was assessed and filtered using Gubbins v2.4.1 (65). Constant sites from the reference genome were obtained using the script iqtree-calc_const_sites.sh (https://github.com/MDU-PHL/mdu-tools/blob/master/bin/iqtree-calc_const_sites.sh). A maximum-likelihood (ML) phylogeny was inferred using IQ-TREE v2.1.0 (66) with parameters set to a GTR+G4 nucleotide substitution model with constant sites and ultrafast bootstrapping (67) to 1000 replicates. The resulting ML tree was then visualised using *ggtree* v2.3.3 R package (68). Population clades were defined using rhierBAPS v1.1.3 (69) to one level on an alignment of 2,957 core SNPS of 1,295 isolates in the ST10 *S.* Dublin lineage.

### In silico prediction of resistance determinants and plasmid replicons

To screen for quinolone point mutations and acquired AMR determinants, ARIBA and AbriTAMR v0.2.2 (https://github.com/MDU-PHL/abritamr) were used, respectively. The AMR database used for this study was the NCBI AMRFinderPlus database v2020-01-22.1. Each isolate’s susceptibility profile was classified as either susceptible; resistant to 1-2 antimicrobials, or multidrug resistant (MDR). Heavy metal and biocide determinants were identified by screening the AMRFinder database v2020-07-16.2 with AMRFinder v3.8.4 (70). Replicon types were determined using ABRicate v1.0.1 (https://github.com/tseemann/abricate) at a threshold of 80% nucleotide identity and 95% coverage, and screening against the Plasmidfinder database (71). A strict threshold was applied to ensure the detection of only unambiguous replicon types. To establish the most common resistant profiles, co-occurrence matrices of resistant determinants were constructed in R using *igraph* v1.2.4.1 (72).

### Virulome analysis

Virulence genes were identified using ABRicate at 90% nucleotide identity/coverage against the virulence factor database (vfdb) v2018-03-18 (73). Databases from previous studies (16,27,35,74–78) were also interrogated and collated. This included a database containing additional virulent determinants and *Salmonella* pathogenicity islands (SPIs) previously implicated with *S*. Dublin invasome (35,41). Using ABRicate, the presence of SPIs -1, -2, -6 and -19 were identified at 80% ID and 10% COV (Supplementary Table 14). ABRicate was also used to screen the manually created SPI determinant database for genes relevant to SPI-1, -2, -6 and -19 with a threshold of 90% identity/coverage (Supplementary Table 14). To quantify invasiveness, index scores were calculated *in silico* for each isolate using Wheeler *et al*.’s tool developed in 2018 for inferring the likelihood of invasiveness (http://www.github.com/UCanCompBio/invasive_Salmonella), with a higher score for an isolate deemed more invasive (38). Prior to running the tool, a set of reference genomes consisting of invasive and non-invasive *Salmonella* serovars were used as a validation set (Supplementary Table 8). The validation results were then visualised using *ggtree*.

### Bayesian phylodynamic analysis

A subset of 660 *S*. Dublin genomes from the main cluster was used for BEAST v1.10.4 (79) analysis. This subset comprised all Australian genomes and a proportion of international *S*. Dublin genomes. The sampling strategy for selecting representative international genomes involved capturing all geographical regions and sources harbouring distinct susceptibility profiles. When two genomes demonstrated the same AMR profile and were from the same region and source, the oldest genome was preferred. Recombination sites were filtered using Gubbins. A core genome SNP alignment of these genomes was created using snippy-core v4.6.0 and then a ML phylogeny was inferred as above. Temporal signal was assessed using TempEst v1.5.3 (80) by assessing the best-fitting root using the heuristic residual mean squared method, with an R^2^ of 0.31. The core alignment was then used to create an xml file in BEAUti v1.10.4 using the GTR+G4 substitution model. To identify the most appropriate model for our dataset, a combination of using either a “relaxed lognormal” or a “strict” molecular clock with either a “coalescent constant” or a “coalescent exponential” population prior was tested. For all models, the Markov chain Monte Carlo chain length was set to 300,000,000 and sampling was set to every 20,000 trees with the marginal likelihood estimation set to “generalised stepping stone (GSS)” to assess the statistical support for models and evaluate the best-fit model. The analysis was repeated in triplicates with and without tip dates to ensure there was no temporal bias. Bayesian evaluation of temporal signal (BETS) was used to determine whether a temporal signal was present (81). To optimise computational resource and time, BEAST v1.10.4 was used with BEAGLE v3.0.2 (82). A Maximum Clade Credibility (MCC) tree from the highest supported model using GSS was extracted using TreeAnnotator v1.10.4 and median heights. The resulting MCC tree was then visualised using *ggtree*.

### Long read sequencing

The Oxford Nanopore Technologies MinION platform was used to obtain complete genomes of six Australian *S*. Dublin genomes (Supplementary Table 13). These genomes, comprised of ST10 lineage clade 1 (AUSMDU00056868 and AUSMDU00035676,) and ST74 (AUSMDU00016676 and AUSMDU00021231), were selected based on their AMR profile, virulome and index scores. To prepare new DNA libraries for long read sequencing, genomes were freshly re-cultured. DNA was extracted using the Virus/Pathogen DSP midi kit on the QiaSymphony SP instrument. Sequencing libraries were prepared using a Ligation sequencing kit (SQK-LSK109) with native barcode expansion and sequenced for 48 hours on a GridION Mk2 instrument using a R9.4 flowcell. WGS was performed using the Illumina NextSeq 500/550 platform (Illumina, San Diego, CA). Demultiplexed long reads and corresponding WGS reads were subjected to QC to exclude unreliable reads using fastp v0.20.1 (83) and Filtlong v0.2.1 (https://github.com/rrwick/Filtlong), respectively. Long reads were assembled using Trycycler v0.4.1 (84). The final sequence was polished using Medaka v1.4.3 (https://github.com/nanoporetech/medaka). To ensure small plasmids were not excluded during the initial QC step, hybrid assemblies using both long and WGS reads were generated Unicycler v0.4.4 (85). Trycycler assemblies were checked against Unicycler assemblies with Bandage v0.8.1 (86). The final genome assemblies were confirmed as *S*. Dublin using SISTR and annotated using both Prokka v1.14.6 (63) for consistency with the draft genome assemblies and Bakta v1.10.1 (87) which provides for more detailed annotations (Supplementary Table 13). Both Prokka and Bakta annotations were in agreement for AMR, HMR and virulence genes, with Bakta annotating between 3-7 additional coding sequences (CDS) which were largely ‘hypothetical protein’.

Chromosomal assemblies were aligned to complete genomes of known *S*. Dublin references CT_02021853 (CP001144.1, without Vi capsule) and CFSAN000518 (CP074226.1, with Vi capsule) and visualised using Blast Ring Image Generator (BRIG) v0.95 (88). Plasmid assemblies were screened against the PlasmidFinder database (71) and mob-typer v3.1.0 (89) to identify replicon and relaxase types, respectively. Plasmid profiles were subjected to BLASTn against previously identified IncX1/IncFII(S) virulence plasmids reported in *S.* Dublin and IncN plasmids associated with serovars of *Salmonella* (Supplementary Table 5). Genome comparisons of plasmids demonstrating the top four highest BLASTn max scores were visualised using BRIG. Bakta, PlasmidFinder and ISFinder (90) were used to annotate the regions of interest, while Easyfig v2.2.2 (91) was used to compare regions of the genome. Regions of the genome were visualised using the R package *gggenomes* v0.9.12.9000 (https://github.com/thackl/gggenomes).

### Human cell line infections

Human macrophage (THP-1) (ATCC® TIB-202™) and colonic epithelial (HT-29) (ATCC® HTB-38™) cell lines were maintained in Roswell Park Memorial Institute (RPMI) 1640 media + 200 mM GlutaMAX (Life Technologies) supplemented with 10% v/v Fetal Bovine Serum (FBS; Bovogen) and grown in a humidified 5% CO2 37°C incubator. THP-1 monocytes were differentiated into macrophages with 50 ng/mL phorbol 12-myristiate-12 acetate (PMA, Sigma-Aldrich) for 72 hours prior to infection.

Single colonies of each *S*. Dublin isolate were inoculated into 10 mL Luria Bertani (LB) broth and grown overnight at 37°C at 200 rpm, then sub-cultured for 3 hours in the same conditions in 5 mL LB. Colony forming units/mL were measured by absorbance at 600 nM (OD_600_). THP-1 and HT-29 cells were infected at a multiplicity of infection (MOI) of 10 in RPMI in triplicate wells of a 96-well plate and centrifuged at 525 x g for 5 minutes to synchronize bacterial uptake. Infected cells were incubated at 37°C for 30 minutes, then infective media removed and replaced with 100 μg/mL gentamicin in 10% v/v FBS/RPMI to inhibit extracellular bacterial growth. 0 hours post infection (hpi) samples were collected at this time to measure initial bacterial uptake. Cells were then incubated in high dose gentamicin for 1 hour, then replaced with 10 μg/mL gentamicin in 10% v/v FBS/RPMI for the remainder of the infection. For THP-1 cells, samples were collected at time 0-, 3-, 9- and 24-hours post-infection. For HT-29 cells, samples were collected at 0- and 24-hours post-infection.

Cytotoxicity of infection was measured by lactate dehydrogenase (LDH) release by infected cells before lysis for intracellular bacterial enumeration. Briefly, 35 uL of infected (and uninfected) cell culture supernatants were collected from each well and LDH quantified as per manufacturer’s instructions (Promega), with percent cytotoxicity calculated as a proportion of lysed control cells and blank media. Following this, infected cells were washed three times with warm phosphate-buffered saline (PBS) to remove any lingering extracellular bacteria, then lysed in 0.1% Triton X-100/PBS and plated out onto LB agar after serial dilution. Bacterial enumeration and cell viability assays were performed over 3 biological replicates, each performed independently, with a new passage of host cells, on a separate day. Within each biological replicate, technical triplicates were performed for each isolate. All statistical analyses were performed using Prism software (GraphPad Software v9.0) and determined by nested students t-test, which allowed for inclusion of individual isolates in sub-groups of each sequence type (ST10 vs ST74).

### Measurement of cellular responses

THP-1 cells were infected at an MOI of 10 with a representative isolate from each of the ST10 and ST74 lineages as described above, then lysed in radio-immunoprecipitation (RIPA) buffer (20 mM Tris pH 8.0, 0.5 mM EDTA, 150 mM NaCl, 1% v/v Triton X-100, 0.5% w/v sodium deoxycholate, 0.1% w/v SDS) supplemented with Complete protease and phosphatase inhibitor cocktails (Roche) at 0-, 3-, 8- and 24 hpi. Total protein concentration was quantified by Bicinchoninic acid (BCA) assay and 15 μg separated on 4-12% Bolt Bis-Tris polyacrylamide gels (Thermo Scientific), transferred to nitrocellulose (iBlot, Thermo Scientific) and blotted for cleaved IL-1β (clone D3A3Z, Cell Signalling). Cell culture supernatants were also collected from 96-well plate infections at 9 hpi and secreted IL-1β measured by DuoSet© Human IL-1β/IL-1F2 ELISA as per manufacturer’s instructions (R&D Systems). A core genome SNP ML phylogeny was constructed on the Australian ST10 lineage to demonstrate the genetic relationship of representative ST10 isolates that underwent *in vitro* phenotypic assays.

### Pangenome analysis

The pangenomes of the isolates used in the macrophages and epithelial infections were analysed using Panaroo v1.3.4 (92) set to default threshold settings, under strict mode and with the removal of invalid genes on the Prokka annotated draft genome assemblies (Supplementary Table 11). A SNP distance matrix was generated using snps-dist v0.8.2 (https://github.com/tseemann/snp-dists) based on the core genome alignment of these isolates, which had been previously used to construct a maximum likelihood (ML) tree.

## DATA AVAILABILITY

Paired end reads of public data were obtained from either the European Nucleotide Archive (ENA) or National Center for Biotechnology Information (NCBI). The Australian short read data and complete genomes for the four Australian *S.* Dublin isolates are available at the NCBI Sequence Read Archive under BioProject PRJNA319593. Details of all 1,303 genomes included in the study are available in Supplementary Tables 1-14. These supplementary tables also includes *i*) the resistance, virulence, Vi and *fliC* profiles of all 1,303 genomes; *ii*) the genbank accession numbers of all reference genomes, virulence determinants and IncN plasmids used in this study, *iii*) genomic annotations of putative novel plasmids, and *iv*) all raw CFU, cytotoxicity and ELISA values from the phenotypic analyses. The interactive tree is available in Microreact under https://microreact.org/project/sdublinpaper).

## CODE AVAILABILITY

The machine learning tool developed by Wheeler *et al*. for predicting the likelihood of invasiveness is available in a public Github depository http://www.github.com/UCanCompBio/invasive_Salmonella.

## ACKNOWLEDGEMENTS

We thank all staff members of the Enterics and MDDI section at the Microbiological Diagnostic Unit Public Health Laboratory (MDU PHL) for their technical assistance. We would also like to thank Jake Lacey for his assistance uploading the reads to NCBI. The BEAST analysis for this paper was undertaken using the LIEF HPC-GPGPU Facility hosted at the University of Melbourne. This Facility was established with the assistance of LIEF Grant LE170100200.

## Contributions

Study design and oversight: CMS, RA, DAW, JSP, DJI. Isolate collection and metadata: MV, PA, SAB, BJP. Performed wet lab experiments: RA, JSP. Data analysis: CMS, RA, JSP, DJI. Manuscript writing: CMS, RA, DAW, JSP, DJI. All authors contributed to manuscript editing.

