## Supplementary Material for "Distinct adaptation and epidemiological success of different genotypes within *Salmonella enterica* serovar Dublin"

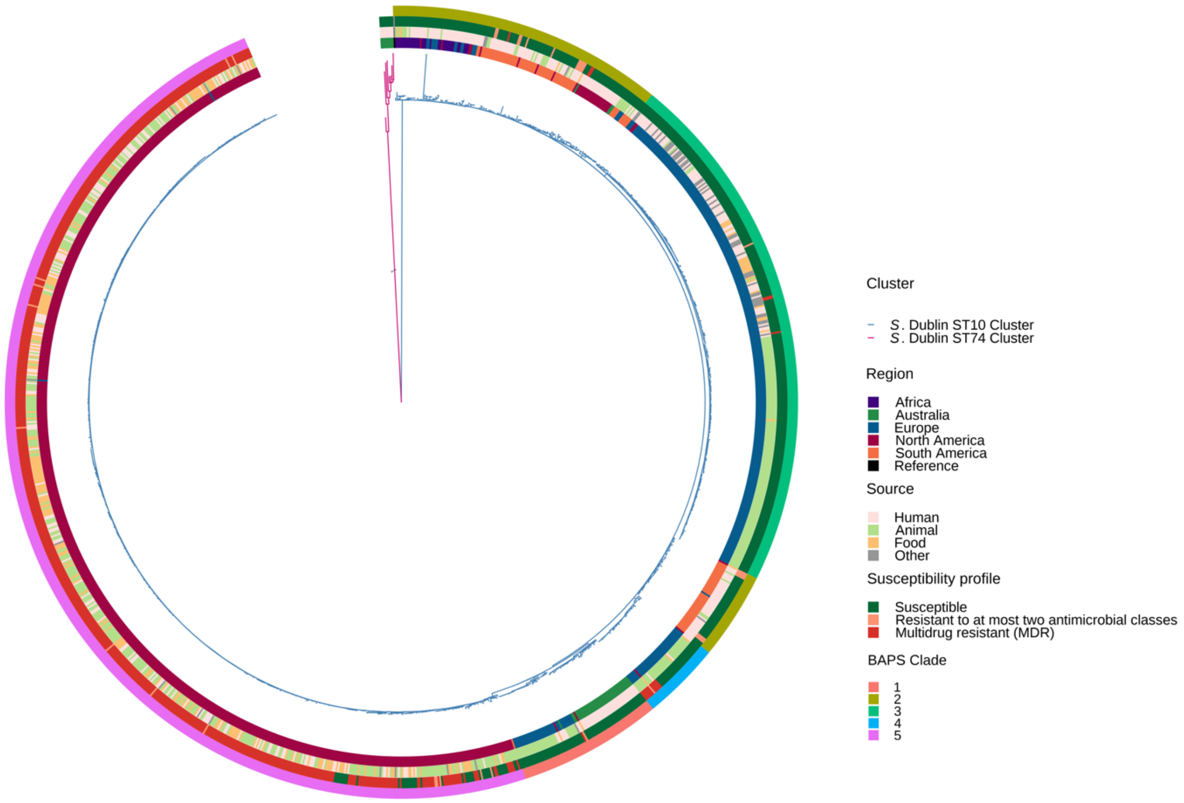


**Supplementary Figure 1. Phylogeny of 1,303 *S.* Dublin genomes.**

Population structure of *S.* Dublin genomes. A) Mid-point rooted maximum likelihood phylogeny inferred from 5,937 SNPs. The branches are coloured by lineage with ST74 in pink and ST10 lineage in blue. The coloured boxes on the tree are the clades identified in the ST10 lineage using the Bayesian Analysis of Population Structure (BAPS). The tips of the tree are coloured by ST. The innermost coloured ring indicates geographical region of collection, the middle ring indicates genome source followed by the AMR susceptibility profile, while the outer ring indicates the BAPs cluster.

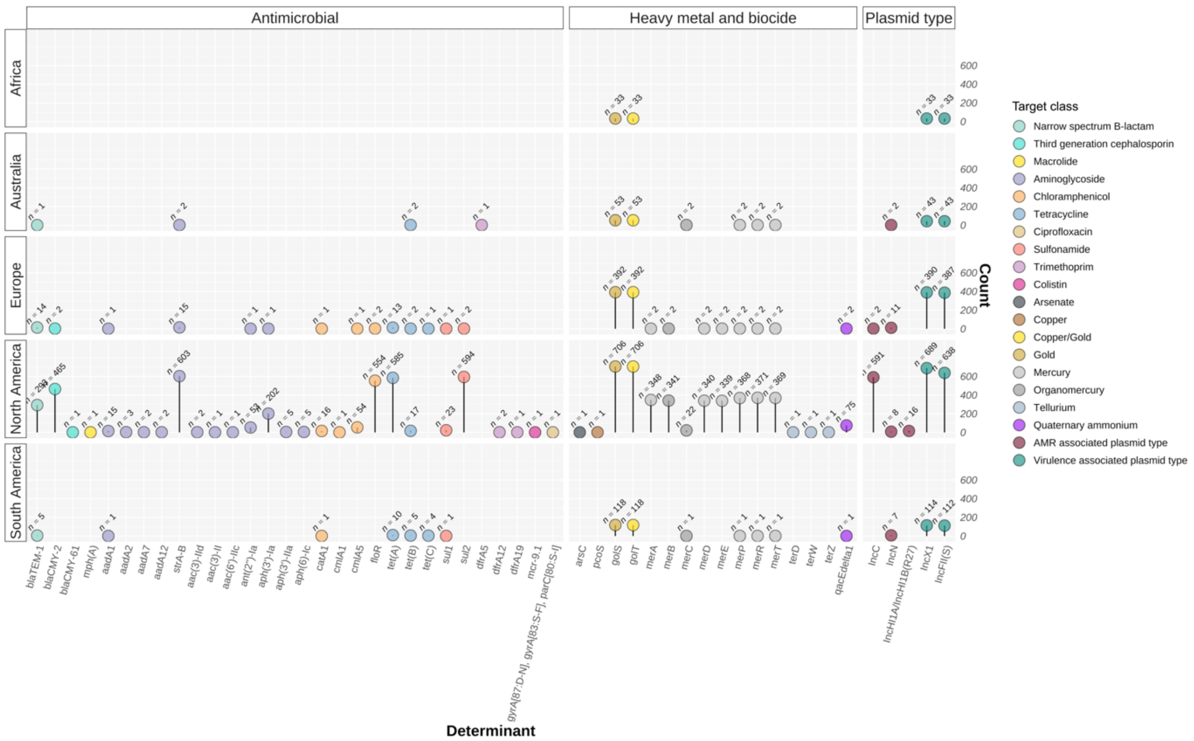


**Supplementary Figure 2. Resistant genotypic profiles *S.* Dublin genomes.**

Distribution of resistant determinants for each region. The number of genomes associated with AMR, heavy metal and biocide determinants and selected plasmid replicons. Each heatmap is coloured according to the target determinant class.

AMR, Antimicrobial resistance.

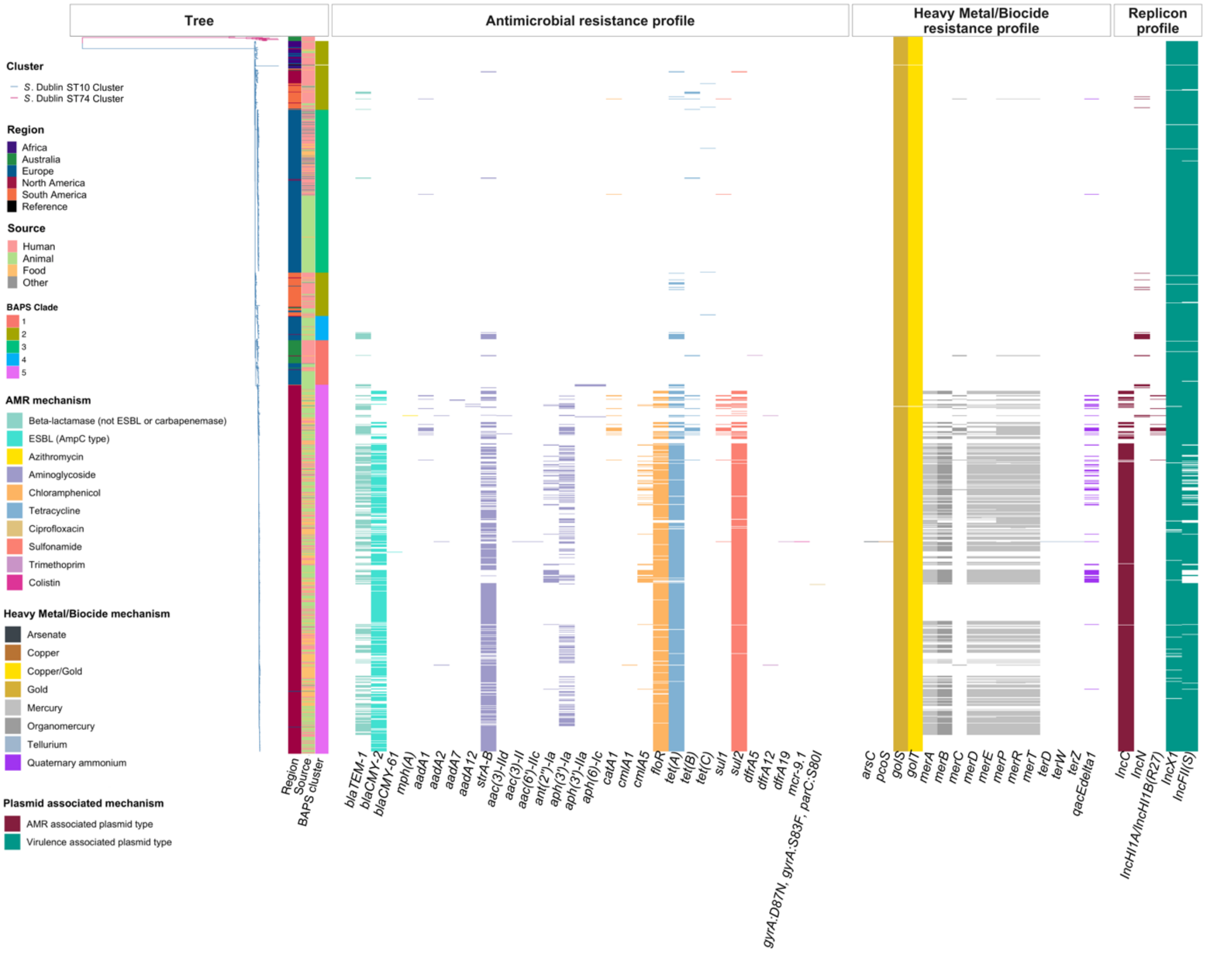


**Supplementary Figure 3. Distribution of resistance determinants in 1,303 *S.* Dublin genomes**

The distribution of antimicrobial resistance determinants, heavy metal resistance determinants, biocide resistance determinants, and selected plasmid replicons detected are shown to the right of the ML tree. Each heatmap is coloured according to the region, the source, BAPs cluster, the target resistance determinant, and the plasmid mechanism.

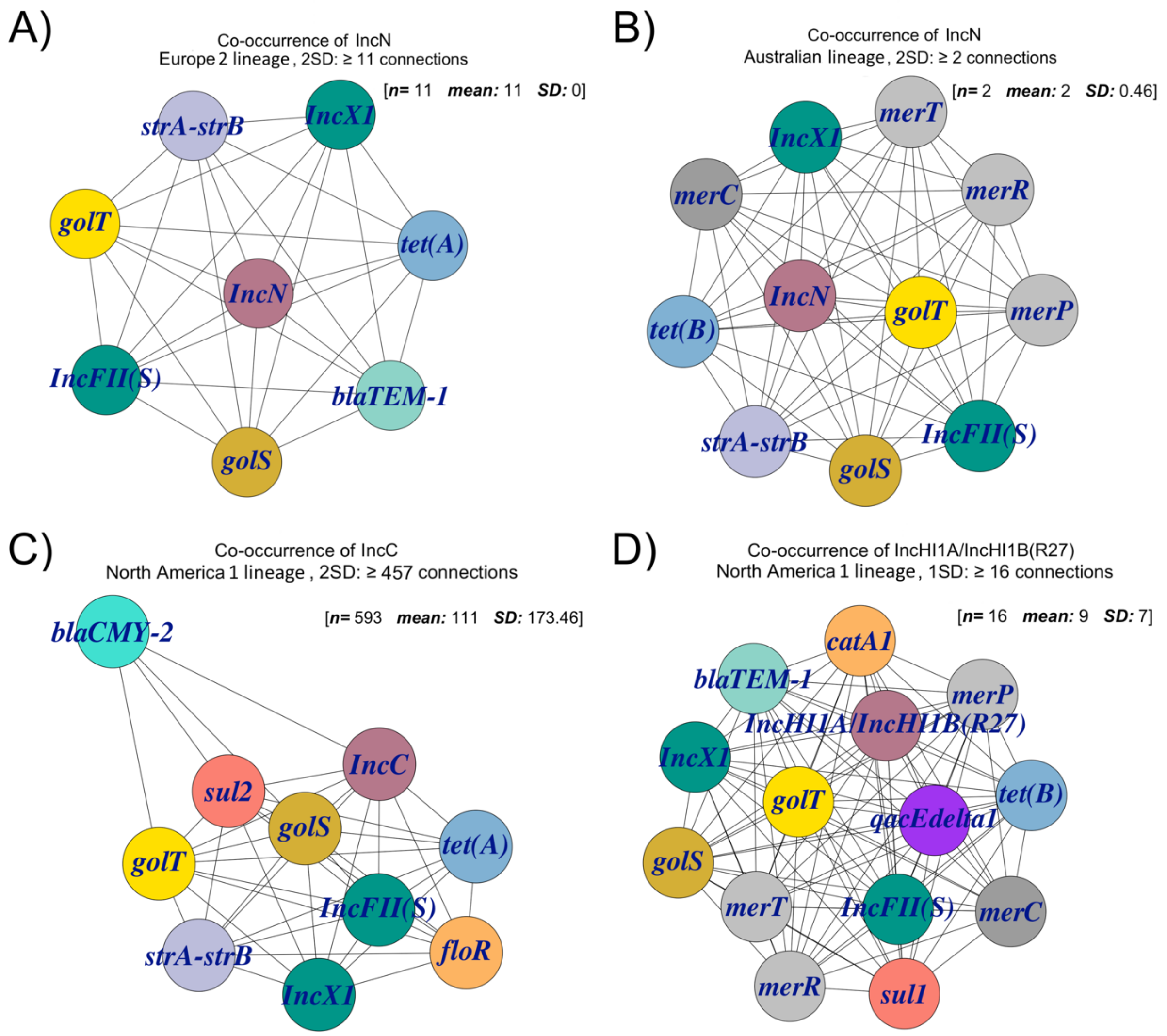


**Supplementary Figure 4. Co-occurrence networks of IncN and IncC resistant profiles**

The most common co-occuring resistant profiles observed for IncN and IncC plasmid types according to region. A) Co-occurrence networks of European AMR genomes harbouring IncN. B) Co-occurrence networks of Australian AMR genomes harbouring IncN. C) Co-occurrence networks of *S.* Dublin genomes from North America harbouring IncC. D) Co-occurrence networks of *S.* Dublin genomes from North America harbouring IncHI1A/IncHI1B(R27). The nodes represent the AMR determinants coloured according to its target determinant class, while the frequency of gene co-occurrence under a given standard deviation is represented by the edges.

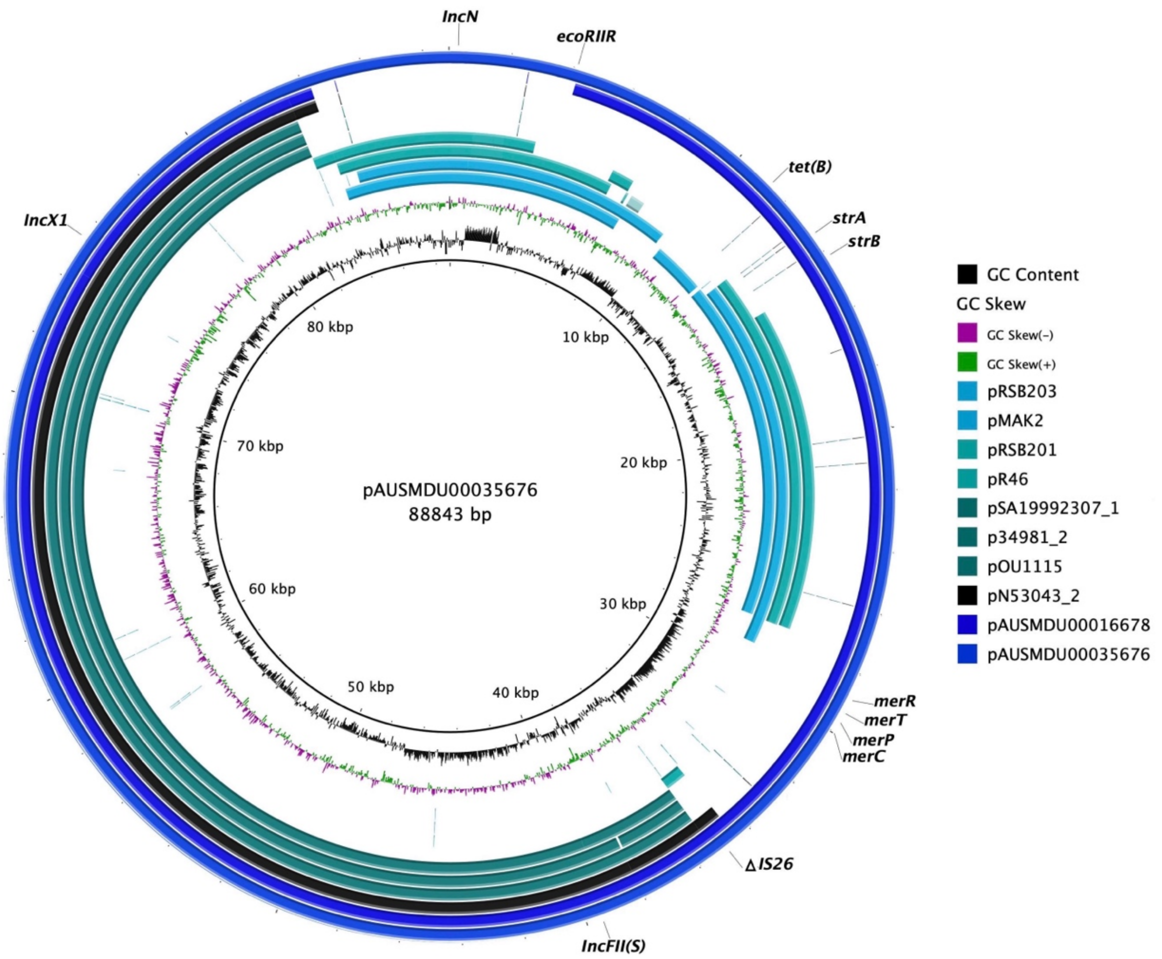


**Supplementary Figure 5. Comparison of plasmids from antimicrobial resistant AUSMDU00035676 and AUSMDU00016678.**

BRIG plots showing sequence homology between complete plasmid genomes from two Australian isolates AUSMDU00035676 (Biosample SAMN40405412) and AUSMDU00016678 (Biosample SAMN18617292). pAUSMDU00035676, a closed reference plasmid (88,843 bp), formed the outermost ring of the plot, while AUSMDU00016678 formed the subsequent ring. The three plasmid replicons, AMR and HMR determinants detected in the plasmid are shown. Comparison plasmids include four IncN plasmids from serovars of *Salmonella* (pSRB203 (JN102342.1) and pSRB201 (JN102341.1) from *S. enterica*, pR46 (AY046276.1) from *S.* Typhimurium and pMAK2 (AB366441.1) from *S.* Dublin, and four virulence plasmids from *S.* Dublin, pSA19992307_1 (CP030208.1), p34981_2 (CP032392.1), pOU1115 (DQ115388.2) and pN53043_2 (CP032386.1).

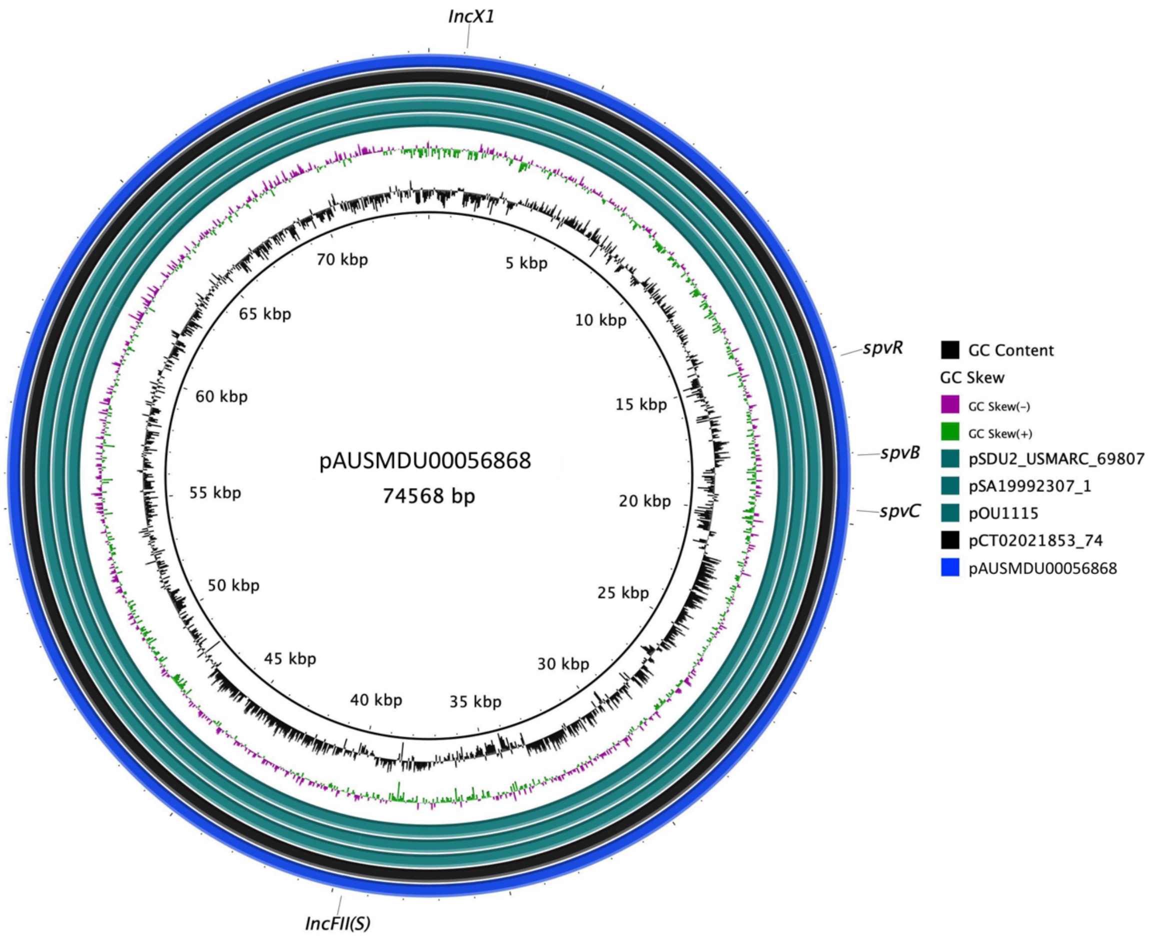


**Supplementary Figure 6. Plasmid assembly of a virulence plasmid identified in an *S.* Dublin ST10 isolate mapped to reference IncX1/IncFII(S) virulence plasmids.**

Plasmid assembly of AUSMDU00056868 (74,568bp), a fully susceptible ST10 isolate, aligned to pCT02021853_74 (CP001143.1, *S.* Dublin), pOU1115, pSA19992307_1, and pSDU2_USMARC_69807 (CP032381.1, *S.* Dublin).

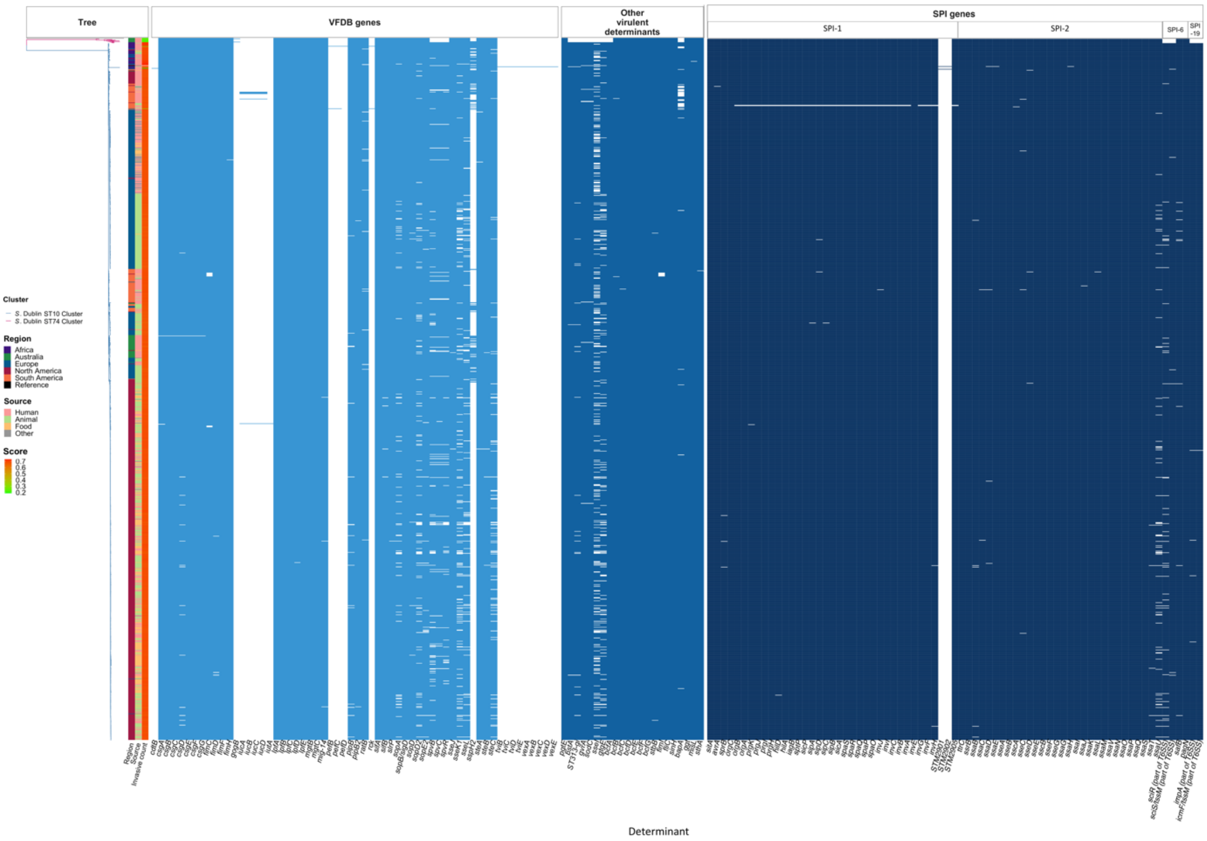


**Supplementary Figure 7. Population structure of *S.* Dublin genomes with genotypic invasive profiles.**

ML tree of the 1,303 is shown on the left. The invasive index score for each isolate is shown on the right of tree. The presences of virulence determinants detected in individual genomes at a 90% coverage and 90% identity threshold are then shown to the right in a heatmap. These determinants are grouped by if they are from the VFDB, were curated from the literature as putative virulence determinants and SPI related genes. The T6SS region in SPI-6 includes four genes and intergenic region and the T6SS region in SPI-19 includes two genes and the intergenic region.

SPI, *Salmonella* pathogenicity island; VFDB, Virulence Factor Database.

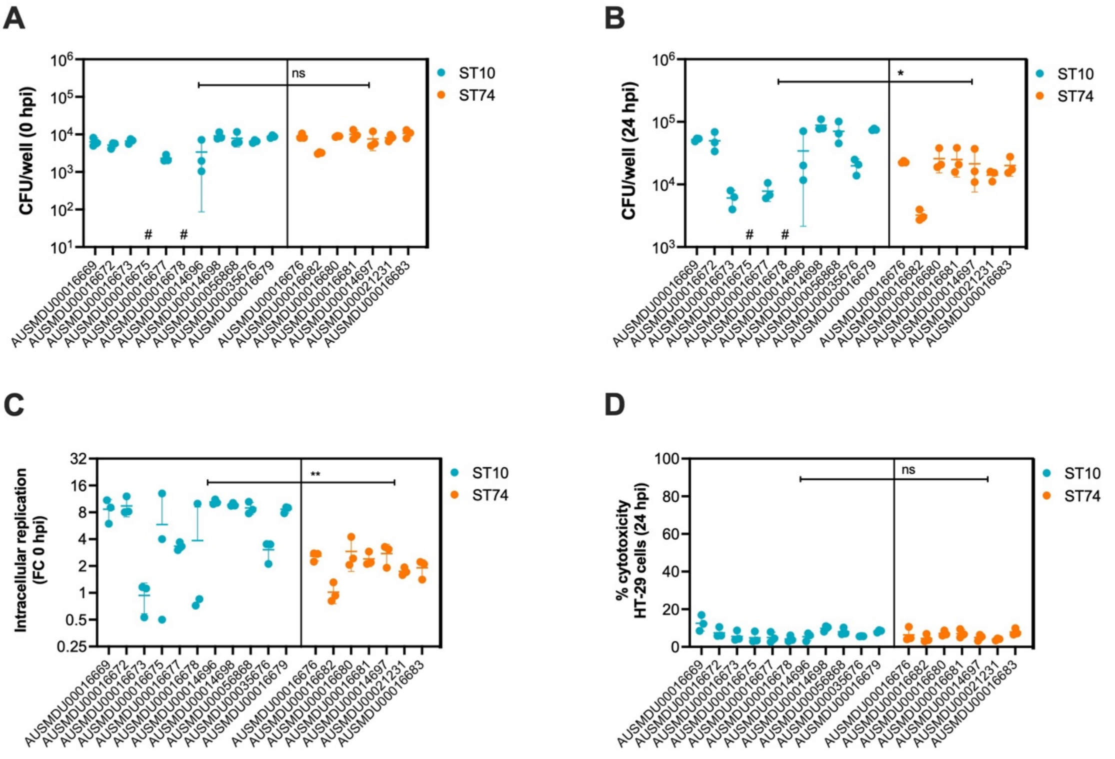


**Supplementary Figure 8. Replication dynamics and host cell cytotoxicity induced by *S.* Dublin ST74 and ST10 lineage isolates in HT-29 cells.**

**A, B)** HT-29 (human intestinal epithelial) cells were infected at MOI:10 with representative *S.* Dublin ST74 or ST10 lineage isolates (Supplementary Table 10). HT-29 cells were lysed, and intracellular bacteria enumerated as CFU/well at the start of infection (0 hpi) and 24 hpi. Each dot represents CFU/well of a biological replicate (performed in technical triplicate), with error bars indicating ± 1 standard deviation of $n$ = 3 biological replicates. **C)** Each dot represents fold change in CFU/well in HT-29 cells (compared to time = 0) of biological replicate (performed in technical duplicate), with error bars indicating ± 1 standard deviation of $n$ = 3 biological replicates. **D)** Cell supernatants were assayed for LDH as a measure of cytopathic effects of infection, and % cytotoxicity was calculated by comparison with 100% lysed uninfected control cells. Samples were collected from HT-29 cells at 24 hpi. Each point represents the % cytotoxicity of a biological replicate (performed in technical triplicate), with error bars indicating ± 1 standard deviation of $n$ = 3 biological replicates. Statistical significance was determined by nested t-test. MOI, multiplicity of infection, hpi, hours post-infection, CFU, colony-forming units, LDH, lactate dehydrogenase, # represents bacterial counts that were below the limit of detection.

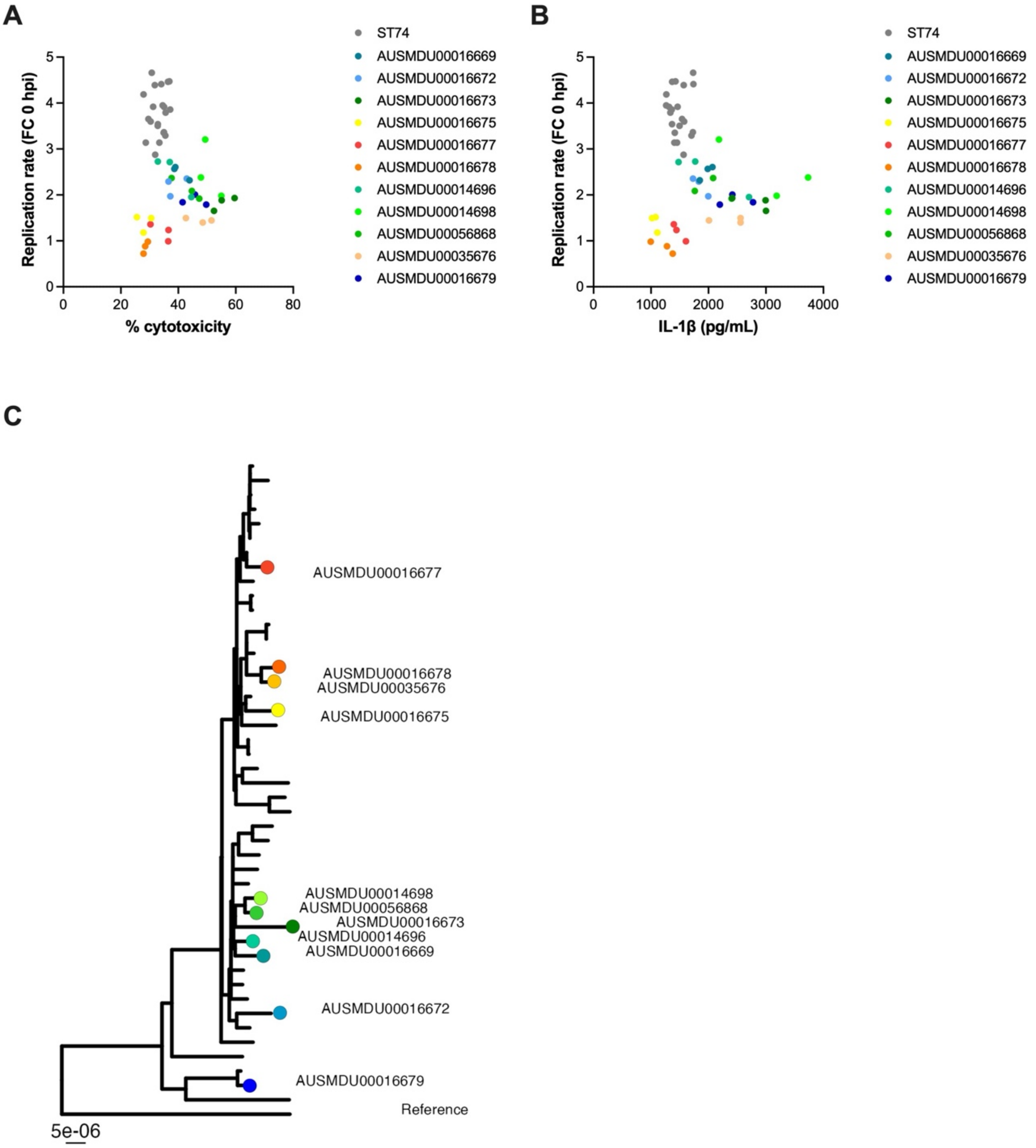


**Supplementary Figure 9. Dot plots comparing bacterial replication and cell death induced by *S.* Dublin ST74 and ST10 lineage isolates in THP-1 cells. (A)** Replication and % cytotoxicity data and **(B)** replication and IL-1β release data were compared on a scatter plot for all *S.* Dublin isolates assayed to show the variation in phenotypic characteristics within the ST10 lineage. Each dot represents one biological replicate (of three) and are coloured according to isolate shown in **(C).**

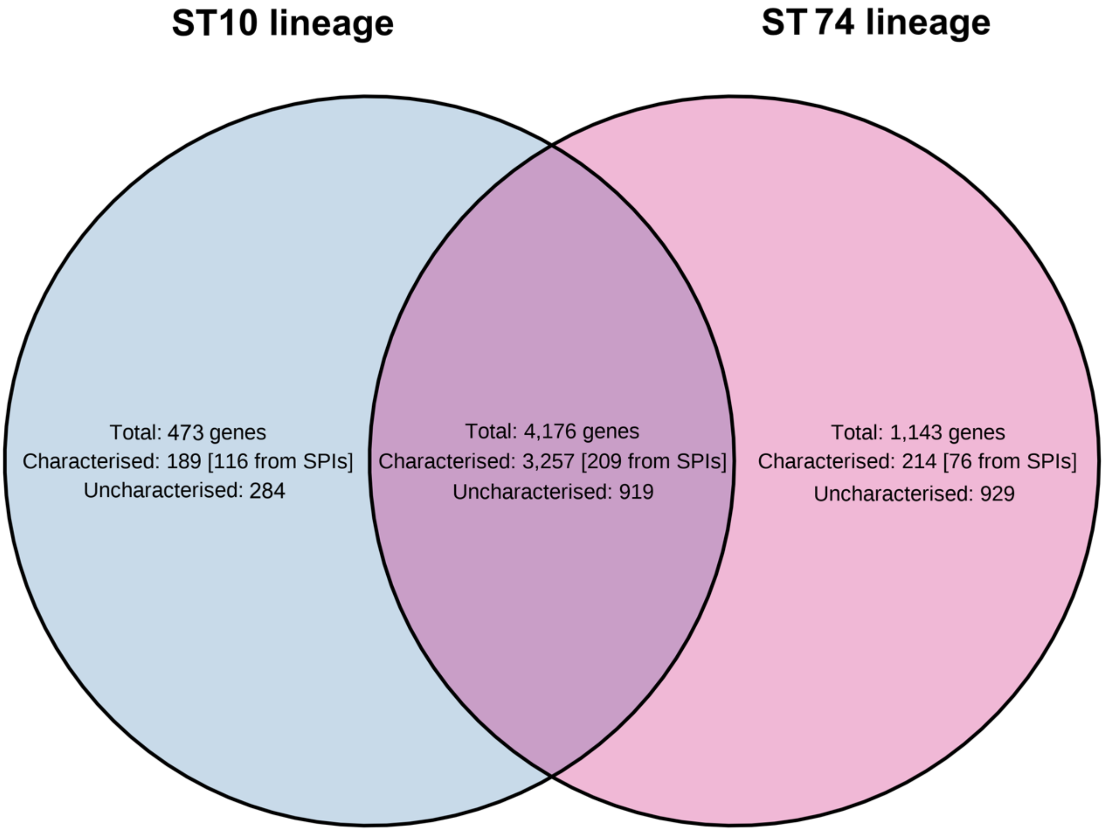


**Supplementary Figure 10: Pangenome visualisation of gene content of ST10 and ST74 lineages**

Venn diagram of gene content of two *S.* Dublin lineages, ST74 (*n* = 7) and ST10 (clade 1) (n = 10).

**Supplementary Table 1. Metadata with genotypic resistance profiles of 1,303 *S*. Dublin genomes.**

**Supplementary Table 2. Distribution of genomes according to source and region.**

**Supplementary Table 3. Distribution of genomes according to sequence type and region.**

**Supplementary Table 4. *fliC* and Vi profiles of 1,303 S. Dublin isolates in study.**

**Supplementary Table 5. Sequence similarity to reference plasmids.**

**Supplementary Table 6. Genome annotations of plasmid AUSMDU00035676**

**Supplementary Table 7. Genome annotations of plasmid AUSMDU00056868**

**Supplementary Table 8. Validation of invasive index prediction tool.**

**Supplementary Table 9. Metadata with virulome profiles of 1,303 *S*. Dublin genomes.**

**Supplementary Table 10. Representative ST10 and ST74 populations used in phenotypic assays.**

**Supplementary Table 11. Results from pangenome analysis of representative isolates from ST10 and ST74 lineages**

**Supplementary Table 12. Single Nucleotide Polymorphisms (SNPs) distance matrix of representative isolates from ST10 and ST74 lineages.**

**Supplementary Table 13. Genomes sent for long read sequencing assemblies.**

**Supplementary Table 14. Database of virulent determinants that have been associated with *S*. Dublin**
